# An improved *Stentor coeruleus* genome and time-resolved transcriptomics link cyclic nucleotide-dependent kinase signaling to single-cell habituation

**DOI:** 10.64898/2026.07.23.740246

**Authors:** Yue Miao, Danielle Khost, Austen Theroux, Nhi Doan, Tejas Ramdas, Samuel J. Gershman

## Abstract

The giant ciliate *Stentor coeruleus* is a single cell capable of modifying its behavior through experience. When repeatedly disturbed, it undergoes habituation, a simple and widely conserved form of non-associative learning, despite lacking a nervous system. How a single cell achieves such behavioral plasticity at the molecular level remains poorly understood. Here, we report a compact, contiguous macronuclear reference genome for *S. coeruleus*, substantially improving genome continuity and annotation quality while resolving two successive rounds of whole-genome duplication and extensive lineage-specific gene-family expansion. Anchoring time-resolved transcriptomics to this reference, we profiled pooled RNA from behaviorally tracked cells collected at defined points during a mechanical stimulation paradigm and identified 341 genes with significant temporal expression dynamics during habituation. The dominant transcriptional signature of habituation was the coordinated, sustained up-regulation of a conserved cyclic nucleotide, calcium, and phosphatase signaling module centered on five cGMP-dependent protein kinase (PKG) paralogs together with a voltage-gated calcium channel and a protein phosphatase 2A regulatory subunit, representing conserved intracellular components of the neuronal long-term depression (LTD) pathway. By integrating an improved genome with time-resolved transcriptomics of habituation, this work provides a community resource for *Stentor* and identifies a conserved signaling program as a candidate molecular framework underlying single-cell habituation.

## 1 Introduction

The capacity to modify behavior with experience is often regarded as a hallmark of animals with nervous systems, yet it is also displayed by single cells. The giant heterotrich ciliate *Stentor coeruleus*, a trumpet-shaped cell that can reach a millimeter in length, has been a model for single-cell biology for over a century, valued for its regenerative abilities and for its capacity to alter its behavior with experience ^1,2^. When mechanically disturbed, *S. coeruleus* rapidly contracts toward its holdfast; upon repeated, inconsequential stimulation the probability of contraction progressively declines, an attenuation that satisfies the operational criteria for habituation, a simple and phylogenetically widespread form of non-associative learning ^3,4^. How a single cell, lacking synapses or a nervous system, stores and updates such information remains a fundamental and largely open question.

Early electrophysiological studies localized the relevant plasticity to a decline in the cell’s mechanoreceptor current^5^, but the molecular machinery underlying single-cell habituation has remained largely undefined. Recent functional work has implicated calcium signaling and protein phosphorylation, showing that perturbing these pathways alters the rate and extent of habituation ^6^. Whether habituation is accompanied by coordinated transcriptional remodeling, however, has never been examined systematically. Addressing this question requires both a well-controlled behavioral paradigm and a high-quality reference genome to support genome-wide transcriptomic analyses.

Such a reference genome has been lacking. Beyond improving gene annotation and transcriptomic analyses, a high-quality genome assembly provides the foundation for reconstructing genome evolution, including whole-genome duplication (WGD) and gene-family evolution, processes that have generated genetic novelty and functional diversification across eukaryotes ^7,8^. Ciliates maintain a separate, highly polyploid, transcriptionally active macronucleus, and ancient WGDs have profoundly shaped genome architecture and gene content in *Paramecium* ^9,10^. Whether similar evolutionary processes have shaped the *Stentor* genome has remained unclear. The first *S. coeruleus* genome assembly revealed unusual genomic features, including exceptionally short introns and macronuclear ploidy that scales with cell size, but remained fragmented ^11^. Moreover, its duplication history has remained contentious: the original study found no evidence for whole-genome duplication, whereas a recent reanalysis inferred an ancient WGD event^12^. Resolving this evolutionary history while establishing a robust foundation for genome-anchored functional studies therefore requires a more complete and contiguous reference genome.

Here we generate a compact, high-quality assembly of the *S. coeruleus* macronuclear genome and use it to address both questions. We first resolve the species’ contested evolutionary history, finding evidence for two successive rounds of whole-genome duplication and extensive lineage-specific gene-family expansion. We then anchor time-resolved transcriptomics to this reference to characterize gene-expression dynamics across a defined series of mechanical stimuli, and identify a coordinated, sustained transcriptional program, centered on cGMP-dependent protein kinase and associated calcium-signaling components, that accompanies habituation. Together, these genomic and transcriptomic resources establish a community reference genome for *Stentor* while identifying conserved signaling pathways as candidate molecular mechanisms underlying single-cell habituation.

## 2 Results

### 2.1 *Stentor coeruleus* genome assemblies vary substantially across assembly strategies

A contiguous and accurate reference genome is essential for investigating genome structure, gene content, and transcriptional regulation in *Stentor coeruleus*. The previously published macronuclear genome, assembled from Illumina short reads, remains highly fragmented, comprising 2,628 contigs with an N50 of only 55 kb ^11^. To generate an improved reference, we sequenced the *S. coeruleus* macronuclear (MAC) genome using PacBio HiFi sequencing, obtaining 9.1 million reads (98.7 Gb; read N50 = 12.6 kb; Table S1), and systematically evaluated multiple assembly strategies.

To identify the most suitable starting assembly for reference-genome construction, we evaluated four assembly strategies: three long-read assemblers, Flye ^13^, HiCanu ^14^, and hifiasm ^15^, together with an iterative assembly approach specifically developed to maximize telomere-to-telomere (T2T) recovery in ciliate genomes ^16^ (Figure S1A). The resulting assemblies differed strikingly in size, contiguity, and completeness (Table 1; Tables S2 and S4). HiCanu generated a 166.28-Mb assembly spanning 4,438 contigs, whereas full-depth hifiasm assembled 909.53 Mb across 17,737 contigs, both exhibiting extensive Benchmarking Universal Single-Copy Orthologs (BUSCO) duplication (80.7% and 97.1%, respectively; Figure 1A). In contrast, Flye produced a compact 63.22-Mb assembly comprising only 263 contigs (N50 = 448 kb) while maintaining high BUSCO completeness (97.1%).

**Figure 1:**
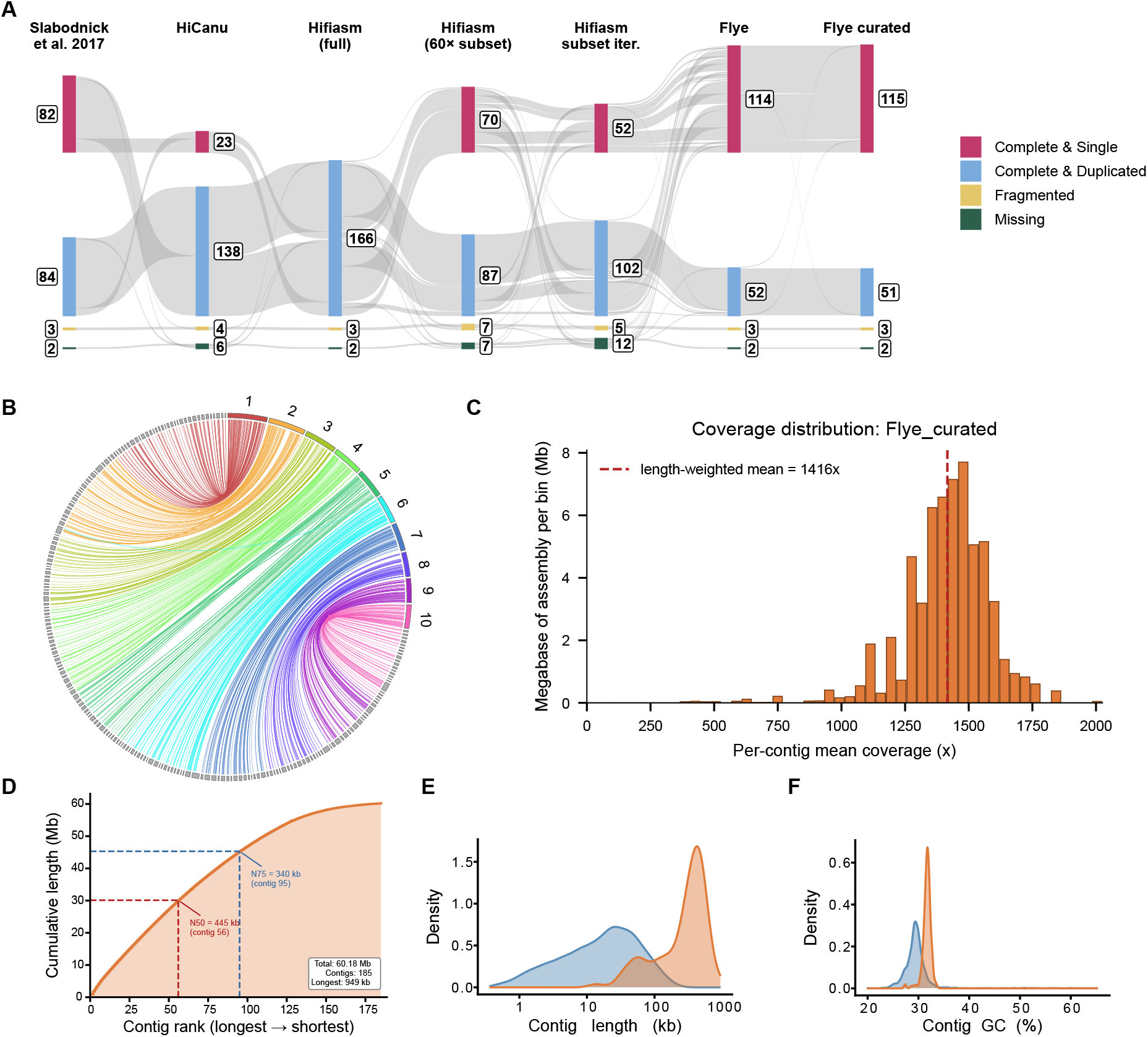
Generation and assessment of the *Stentor coeruleus* reference genome assembly. **(A)** Sankey diagram showing BUSCO category transitions across seven *S. coeruleus* genome assemblies using the alveolata odb10 dataset (*n* = 171). Categories indicate complete single-copy, complete duplicated, fragmented, and missing BUSCOs. Numbers within each box indicate the number of BUSCOs assigned to that category. **(B)** Whole-genome alignment between the curated Flye assembly (SteCoeF v1) and the previously published Illumina assembly. Colored contigs represent SteCoeF v1 and grey contigs represent the 2017 assembly. Links indicate aligned genomic regions and are colored according to the corresponding SteCoeF v1 contig. Contigs are ordered by length, with the ten longest SteCoeF v1 contigs labeled. **(C)** Per-contig mean coverage of the curated Flye assembly; dashed line, length-weighted mean (~1,416×). **(D)** Cumulative contig-length distribution (contigs ordered longest to shortest); dashed lines, N50 (445 kb) and N75 (340 kb). **(E)** Contig-length density and **(F)** contig GC-content density for the Slabodnick et al. (2017) assembly (blue) and the curated Flye assembly (orange).

**Table 1:** Assembly statistics for *Stentor coeruleus* genome assemblies.

| Assembly Statistics | Slabodnick et al. 2017 | Flye | HiCanu | Hifiasm (full) | Hifiasm (60× subset) | Hifiasm subset iter. | Flye curated |
| --- | --- | --- | --- | --- | --- | --- | --- |
| # Contigs | 2,628 | 258 | 4,438 | 17,737 | 1,105 | 382 | 185 |
| Assembly size (Mb) | 77.83 | 63.22 | 166.28 | 909.53 | 89.11 | 86.02 | 60.18 |
| N50 (kb) | 55 | 448 | 43 | 69 | 124 | 378 | 445 |
| Longest contig (Mb) | 0.27 | 1.47 | 1.6 | 4.35 | 2.15 | 1.33 | 0.95 |
| BUSCO (alveolata_odb10, n = 171) | C:97.1%<br>[S:48%,<br>D:49.1%],<br>F:1.8%,<br>M:1.1% | C:97.1%<br>[S:66.7%,<br>D:30.4%],<br>F:1.8%,<br>M:1.1% | C:94.2%<br>[S:13.5%,<br>D:80.7%],<br>F:2.3%,<br>M:3.5% | C:97.1%<br>[S:0%,<br>D:97.1%],<br>F:1.8%,<br>M:1.1% | C:91.8%<br>[S:40.9%,<br>D:50.9%],<br>F:4.1%,<br>M:4.1% | C:90%<br>[S:30.4%,<br>D:59.6%],<br>F:2.9%,<br>M:7.1% | C:97.1%<br>[S:67.3%,<br>D:29.8%],<br>F:1.8%,<br>M:1.1% |
| Inspector Quality Value (QV) | 9.06 | 47.82 | 39.26 | 39.55 | 33.34 | 38.27 | 54.38 |
| Structural errors | 0 | 0 | 0 | 0 | 0 | 7 | 1 |

The iterative assembly strategy substantially improved contiguity relative to the initial subset assemblies, reducing contig number from 1,105 to 382, increasing N50 from 124 kb to 378 kb, and recovering the highest proportion of telomere-capped contigs (59.2%; Table S3). However, despite its superior telomere recovery, the final iterative assembly remained substantially larger than the Flye assembly, exhibited lower BUSCO completeness (90.0%), and introduced multiple structural errors (Table 1; Tables S2 and S3). The marked inflation of the HiCanu and full-depth hifiasm assemblies is consistent with the highly polyploid nature of the *S. coeruleus* MAC genome, in which extensive allelic and paralogous variation can lead assemblers to represent the same locus multiple times. Based on these comparisons, the Flye assembly and the iterative assembly were selected for further curation and contaminant removal, yielding the final assemblies SteCoeF v1 and SteCoeH v1, respectively. Although SteCoeH v1 maximized telomere recovery, it did so at the expense of completeness and structural accuracy, it was retained for independent validation, whereas SteCoeF v1 was selected as the primary reference genome for downstream analyses.

### 2.2 Generation of a compact high-quality reference genome

Starting from the compact Flye assembly, we sequentially removed redundant contigs, corrected local assembly errors, and eliminated contaminating sequences, generating the final reference assembly, SteCoeF v1, comprising 185 contigs spanning 60.18 Mb (Table 1).

Curation substantially improved assembly quality while preserving completeness. Inspector quality value increased from 47.82 to 54.38, and small-scale assembly errors were reduced from 981 to one, the remaining error localizing to an rRNA tandem-repeat region and likely reflecting the repetitive nature of this locus rather than a bona fide misassembly (Table 1; Table S2). BUSCO completeness remained high (97.1%), while Sankey analysis showed that Flye and SteCoeF v1 retained the largest and most stable sets of complete single-copy BUSCOs, in contrast to the extensive duplication observed in HiCanu and hifiasm (Figure 1A). The remaining duplicated BUSCOs in SteCoeF v1 therefore likely reflect genuine biological duplication rather than assembly artifacts.

Whole-genome alignments further supported the accuracy of SteCoeF v1. Multiple short contigs from the 2017 Illumina assembly aligned collinearly along individual SteCoeF v1 contigs (Figure 1B) and SteCoeH v1 contigs (Figure S1B), consistent with consolidation of previously fragmented genomic regions. Alignment between the two HiFi assemblies demonstrated high sequence concordance, with an average nucleotide identity of 99.62% across aligned regions. In addition, 99.1% of SteCoeH v1 assembly aligned to SteCoeF v1, indicating that its larger size primarily reflects redundant representation rather than substantial unique genomic sequence (Table S5; Figure S1C). HiFi read coverage across SteCoeF v1 was unimodal, with a length-weighted mean coverage of approximately 1,416× (Figure 1C), and the assembly exhibited a pronounced shift toward longer contigs and a narrower, slightly higher GC-content distribution than the 2017 assembly (Figures 1D–F). Collectively, these analyses establish SteCoeF v1 as a compact, highly contiguous reference genome for *S. coeruleus* that provides the foundation for all subsequent analyses.

### 2.3 Annotation of the *Stentor coeruleus* genome

With SteCoeF v1 established as the reference genome, we generated a comprehensive structural and functional annotation and compared it with the previously published *S. coeruleus* annotation ^11^ and two congeneric *Stentor* genomes ^12,17^.

Using a ciliate-aware annotation pipeline optimized for the exceptionally short introns of *S. coeruleus* ^17,18^, we identified 32,306 protein-coding genes, slightly fewer than the 34,506 genes reported previously. The annotation retained the characteristic compact gene architecture of *S. coeruleus*: only 18.7% of genes (6,029) contained introns, gene models were predominantly single-exon (mean 1.28 exons per gene; Figure 2A), and intron lengths remained sharply centered at 15 bp (mean 15.6 bp; Figure 2B), consistent with the hallmark tiny-intron architecture previously described for *S. coeruleus* ^11^. Compared with the 2017 annotation, the SteCoeF v1 annotation exhibited highly similar distributions of coding-sequence (CDS) GC content, gene length, exon length (Figures 2C–E), indicating that the improved assembly preserved the underlying gene architecture. Despite the modest reduction in gene number, the SteCoeF v1 annotation was more compact and gene-dense, with a larger fraction of the genome occupied by gene regions (69.1% versus 55.5%) and shorter intergenic distances (mean 560 bp versus 875 bp; Figure 2F; Table S6). CDS GC content was also slightly higher (34.6% versus 32.5%), consistent with the genome-wide GC-content differences between assemblies (Figure 1F). The same architectural features were conserved in *S. pyriformis* and *S. roeselii*, which also exhibited compact, gene-dense genomes with predominantly single-exon gene models and highly similar exon and intron length distributions (Figure S2; Table S6).

**Figure 2:**
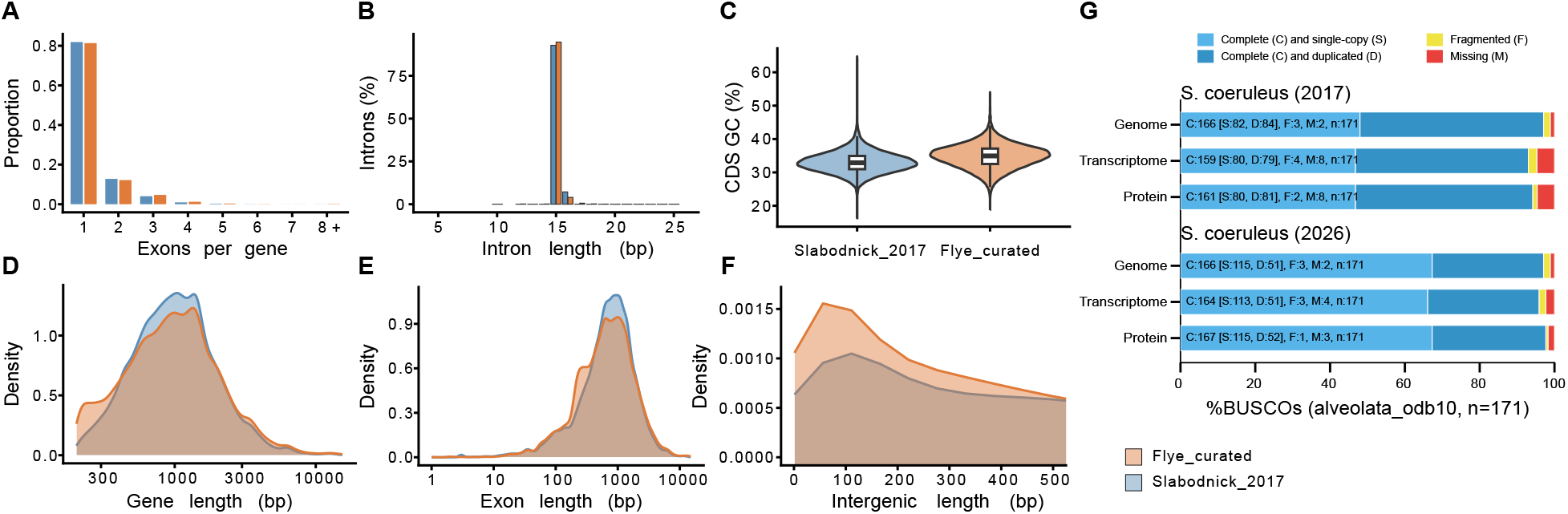
Structural annotation of the *Stentor coeruleus* genome and comparison with the 2017 annotation. Distribution of gene-structure features for the SteCoeF_v1 (curated Flye; orange) and the Slabodnick et al. (2017) annotation (blue): **(A)** number of exons per gene, **(B)** intron length, **(C)** CDS GC content, **(D)** gene length, **(E)** exon length, and **(F)** intergenic length. **(G)** BUSCO completeness (alveolata odb10, *n* = 171) assessed in genome, transcriptome, and protein modes for the 2017 and current SteCoeF_v1 *S. coeruleus* annotations. Bars are colored by BUSCO category (complete and single-copy, complete and duplicated, fragmented, and missing), with corresponding counts indicated for each mode. Numeric gene-structure statistics are provided in Table S6.

We next evaluated annotation completeness in transcriptome and protein modes using BUSCO (Figure 2G; Table S4). In both modes, the SteCoeF v1 annotation recovered more complete orthologs and fewer missing orthologs than the 2017 annotation, extending the improvements in complete single-copy ortholog representation observed in genome mode. These results indicate that the reduction in gene number reflects consolidation of redundant gene models rather than loss of conserved gene content.

Functional annotation integrated homology, orthology, protein-domain, and KEGG evidence. Overall, 93.3% of predicted genes received support from at least one annotation source, and 65.5% were assigned an informative functional description, leaving 34.5% of genes uncharacterized. Gene Ontology (GO) terms were assigned to 15,295 genes (47.3%), and 12,513 genes (38.7%) received KEGG Ortholog assignments (Table S7). This annotated reference genome enabled the whole-genome duplication, gene-family evolution, and transcriptomic analyses presented below.

### 2.4 The *Stentor coeruleus* genome bears signatures of two rounds of whole-genome duplication

Whole-genome duplication (WGD) has profoundly shaped genome evolution across eukaryotes, creating opportunities for gene diversification and the emergence of novel biological functions. Using the high-contiguity SteCoeF v1 assembly, we searched for evidence of ancient WGD through genome-wide collinearity analysis. Self-comparison of the *S. coeruleus* proteome followed by MC-ScanX collinearity detection ^19^ identified 480 intragenomic collinear blocks comprising 7,125 syntenic paralog pairs, with 11,158 genes (34.5% of the predicted proteome) assigned to collinear blocks (Figure 3A; Table S8). Most syntenic paralog pairs linked genes on different contigs (6,988 between-contig versus 137 within-contig pairs; Table S9), indicating that the duplication signal is broadly distributed throughout the genome rather than being confined to local genomic regions.

**Figure 3:**
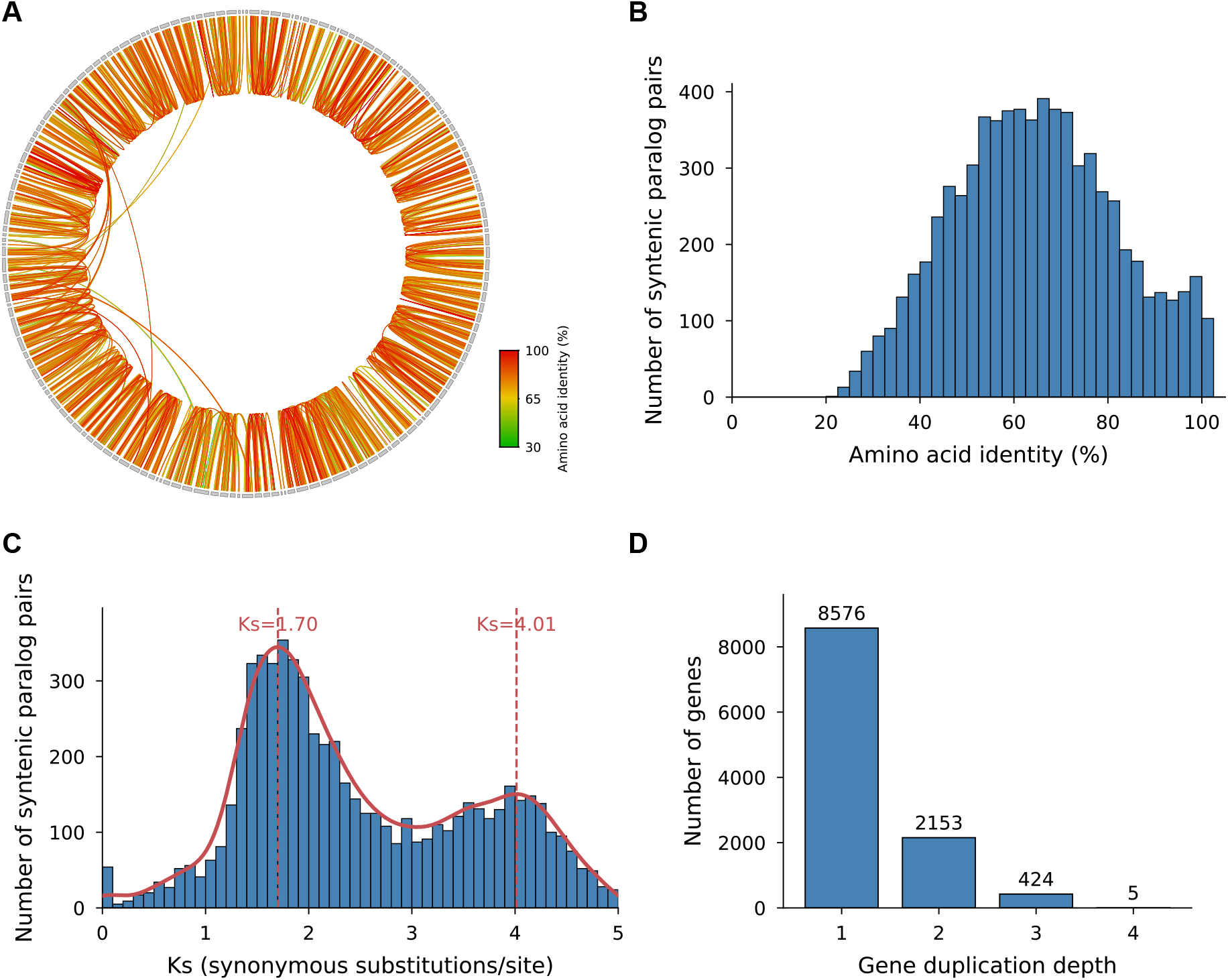
Whole-genome duplication in *Stentor coeruleus*. **(A)** Circos plot of intragenomic collinearity in SteCoeF v1. Contigs are ordered by collinearity-based seriation so that strongly collinear contigs are positioned near one another. Links connect 7,125 syntenic paralog pairs across 480 collinear blocks and are colored by amino-acid identity. **(B)** Distribution of amino-acid identity among the 7,125 syntenic paralog pairs. **(C)** Distribution of synonymous substitution rates (*K_s_*) for syntenic paralog pairs with valid estimates in the range 0 *< K_s_ <* 5 (*n* = 6,392). The red curve indicates a Gaussian kernel-density estimate, and dashed lines mark the two density peaks at *K_s_ ≈* 1.70 and *K_s_ ≈* 4.01. **(D)** Gene duplication depth distribution across collinear genes.

To determine whether these duplicated regions represent ancient biological duplications rather than residual assembly redundancy, we examined amino acid identity among syntenic paralog pairs. Sequence identity was broadly distributed, with a median of 64.3% and values spanning approximately 22–100% (Figure 3B), inconsistent with a genome dominated by residual haplotigs or macronuclear polyploidy artifacts, which would instead produce an enrichment of nearly identical paralogs. We next estimated synonymous substitution rates (*K_s_*) for syntenic paralog pairs using the Yang–Nielsen maximum-likelihood method ^20^. Among 6,392 informative pairs (0 *< K_s_ <* 5), the *K_s_* distribution was clearly bimodal, with density peaks at *K_s_ ≈* 1.70 and *K_s_ ≈* 4.01, separated by a valley at *K_s_ ≈* 3.05 (Figure 3C), consistent with two temporally distinct large-scale duplication events. Gene-duplication depth provided independent support for this inference: among collinear genes, 8,576 exhibited duplication depth 1, 2,153 depth 2, and 424 depth 3 (Figure 3D). Using a conservative threshold that excluded duplication depths represented by fewer than 100 genes, the maximum supported depth was 3, corresponding to four retained copies and therefore two rounds of WGD ^10^. Together, amino acid divergence, the bimodal *K_s_* distribution, and gene-duplication depth provide complementary evidence supporting two successive whole-genome duplication events in *S. coeruleus*.

To place these findings in an evolutionary context, we extended the collinearity analysis to two congeneric species (Table S8). Intragenomic collinearity was abundant in the contiguous *S. coeruleus* assembly but much weaker in the more fragmented *S. pyriformis* and *S. roeselii* assemblies, which yielded only 14 and 1 intragenomic collinear blocks, respectively. The few intragenomic pairs detected in *S. pyriformis* clustered near 100% amino acid identity, consistent with residual assembly redundancy rather than ancient WGD-derived paralogy. In contrast, intergenomic collinearity was readily recovered and was substantially more extensive between *S. coeruleus* and *S. pyriformis* than between *S. coeruleus* and *S. roeselii* (984 versus 262 blocks; Figure S3; Table S8), consistent with their closer evolutionary relationship. These results help reconcile earlier discordant reports: the 2017 short-read *S. coeruleus* assembly did not recover evidence of WGD ^11^, whereas later analyses suggested a duplication signal ^12^. The improved contiguity of SteCoeF v1 reveals extensive genome-wide collinearity and supports a history of two successive rounds of whole-genome duplication.

### 2.5 Lineage-specific gene-family expansion in *Stentor coeruleus*

To examine how the duplicated *S. coeruleus* genome subsequently evolved, we compared gene-family evolution across eight ciliate genomes using OrthoFinder ^21^. Among the three *Stentor* species, 7,833 orthogroups were shared, whereas lineage-specific innovation differed markedly (Figure 4A; Table S10). *Stentor coeruleus* contained 1,218 species-specific orthogroups comprising 5,232 genes, substantially more than *S. roeselii* (546 orthogroups) and *S. pyriformis* (147 orthogroups), indicating greater lineage-specific gene innovation in *S. coeruleus*.

**Figure 4:**
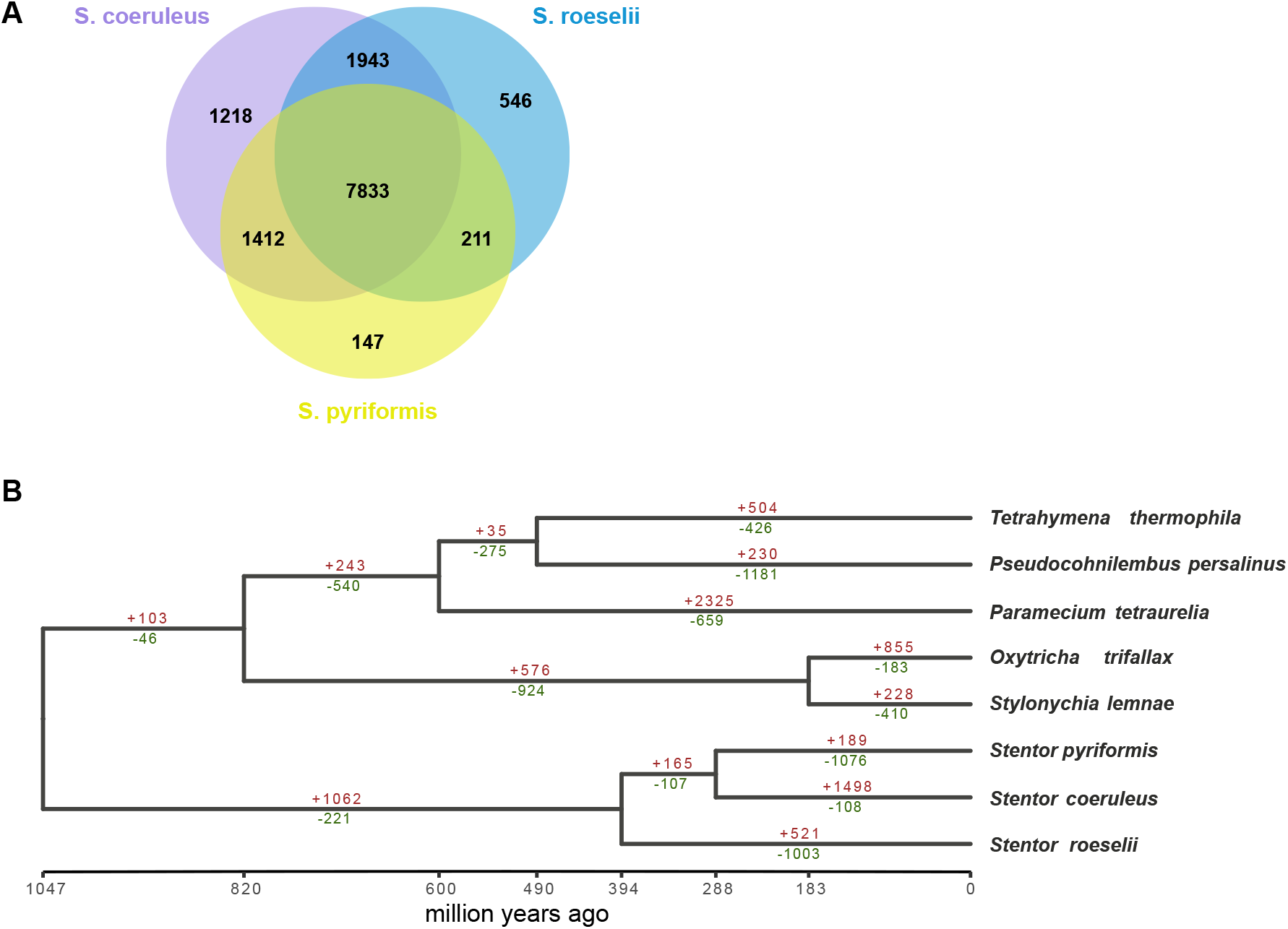
Gene-family evolution across ciliates and lineage-specific expansion in *Stentor coeruleus*. **(A)** Orthogroup sharing among the three *Stentor* species (*S. coeruleus*, *S. pyriformis*, and *S. roeselii*); numbers indicate orthogroup counts in each region. **(B)** Time-calibrated species tree of eight ciliate genomes inferred using OrthoFinder and dated with r8s (axis in millions of years ago) ^23^. Numbers above and below each branch indicate the numbers of gene families inferred by CAFE to have expanded (red, +) or contracted (green, −), respectively, along that lineage.

We next modeled gene-family size evolution using CAFE^22^ on a time-calibrated species tree. In contrast to *S. pyriformis* (189 expanded versus 1,076 contracted families) and *S. roeselii* (521 expanded versus 1,003 contracted families), the terminal *S. coeruleus* lineage exhibited a pronounced excess of gene-family expansion (1,498 expanded versus 108 contracted families; Figure 4B; Table S11). Expansion also predominated in *Paramecium tetraurelia* (2,325 expanded versus 659 contracted families), whose genome has undergone multiple documented WGDs ^9^. These results indicate that the evolutionary history of *S. coeruleus* has been characterized by substantial lineage-specific gene-family expansion.

To characterize the biological functions associated with lineage-specific innovation and gene-family expansion, we performed GO and KEGG enrichment analyses of genes from *S. coeruleus*-specific orthogroups (5,232 genes; Figure S4A; Table S12) and significantly expanded gene families (2,922 genes; Figure S4B; Table S13). The *S. coeruleus*-specific gene set was predominantly enriched for signaling functions, including G-protein-coupled receptor (GPCR) signaling, adenylate and guanylate cyclase activity, protein phosphorylation, and GTPase activity. In contrast, expanded gene families were primarily enriched for transport and cytoskeletal functions, including transmembrane transport, ABC transporters, microtubule organization, tubulin binding, and protein polyglutamylation. Transport-related categories and GPCR-mediated signaling were prominent in both datasets. Although several enriched KEGG pathways carry metazoan-specific names (for example, taste transduction and dopaminergic synapse), the underlying *S. coeruleus* orthologs correspond primarily to GPCRs, ion channels, transporters, and cyclic nucleotide signaling components. These results indicate that lineage-specific evolution in *S. coeruleus* preferentially expanded gene families associated with signaling, transport, and cytoskeletal functions.

### 2.6 Time-resolved transcriptomic profiling of habituation in *Stentor coeruleus*

Habituation in *S. coeruleus* provides a tractable system for investigating how a single cell adapts its behavior in response to repeated experience. To characterize the molecular correlates of habituation, we performed time-resolved RNA sequencing across a standardized mechanical stimulation paradigm. Using an automated behavioral platform ^24^, substrate-anchored *S. coeruleus* cells received calibrated mechanical stimuli at one tap per minute while contraction behavior was quantified from video recordings (Figures 5A and 5B). Independent cell cohorts were collected after 0, 10, 20, 30, 40, 50, and 60 stimuli (Figure 5C). Because only a subset of cells remained anchored throughout the assay, individual cells were tracked continuously, and only those that remained attached and experienced the complete stimulation protocol up to their sampling point were collected for bulk RNA sequencing (Figure 5D). Three biological replicates were obtained for each time point, yielding 21 RNA-seq libraries in total. Across biological replicates, contraction probability progressively declined with repeated stimulation (Figure 5E; Table S14), confirming that the sampled cells collectively span the full behavioral progression from the näıve to the habituated state.

**Figure 5:**
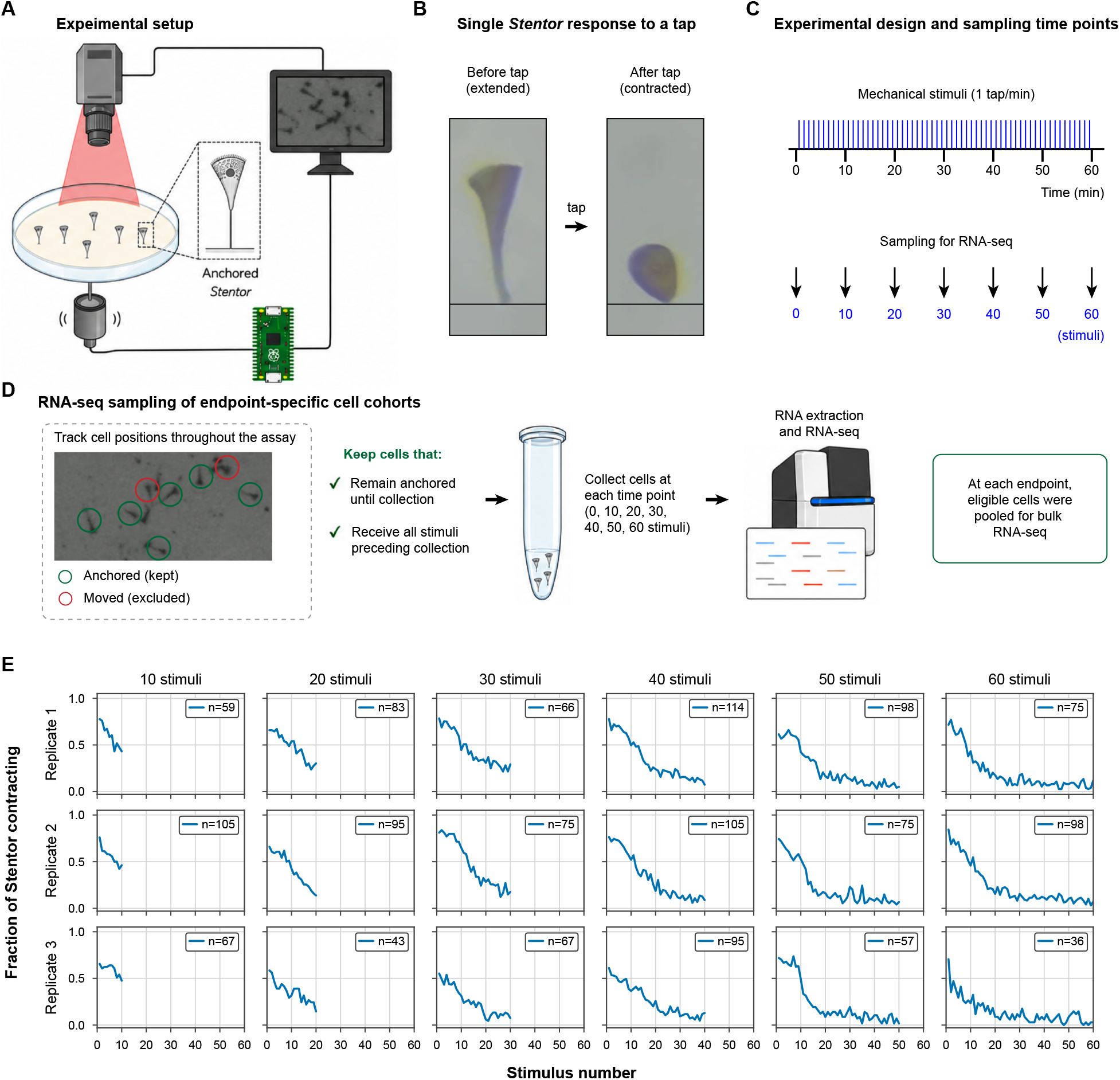
An experimental paradigm for time-resolved transcriptomic profiling of habituation in *Stentor coeruleus*. **(A)** Experimental setup: substrate-anchored *Stentor* cells in a dish are imaged from above while an automated device delivers calibrated mechanical taps. **(B)** A single *Stentor* contracting from the extended to the contracted state in response to a tap. **(C)** Experimental design: stimuli are delivered at one tap per minute, and cells are sampled for RNA-seq after 0, 10, 20, 30, 40, 50, and 60 stimuli. **(D)** RNA-seq sampling strategy: for each independent endpoint cohort, cells that remained anchored throughout the stimulation period up to collection were pooled for bulk RNA sequencing; cells that detached or moved before collection were excluded. **(E)** Behavioral habituation curves for three biological replicates (rows) across the six stimulated endpoints (columns); each curve shows the percentage of cells that contracted as a function of stimulus number. n, number of anchored cells analyzed per replicate.

RNA sequencing generated consistently high-quality transcriptomes across all 21 libraries. Reads aligned to the SteCoeF v1 reference genome using a short-intron-aware HISAT2 configuration, with a mean alignment rate of 97.4% (Figures S5A). After removal of reads overlapping rRNA loci, an average of 93.7% of primary mapped reads was assigned to annotated genes (Figures S5B; Table S15). This high-quality dataset spanning the complete habituation time course provided the basis for subsequent transcriptional analyses.

### 2.7 Transcriptional dynamics of habituation in *Stentor coeruleus*

To identify transcriptional changes accompanying habituation, we applied a likelihood-ratio test across all seven sampling time points, identifying 341 genes with significant temporal expression dynamics (Figure 6A). K-means clustering resolved these genes into four major temporal trajectories (Figure 6B): a transient early-response cluster peaking near 20 stimuli (C1, *n* = 98), a transient late-response cluster peaking near 40 stimuli (C2, *n* = 83), a sustained activation cluster showing progressively increasing expression throughout repeated stimulation (C3, *n* = 118), and a gradually down-regulated cluster (C4, *n* = 42). Functional enrichment analyses for the individual temporal clusters are provided in Table S17. Considering all 341 habituation-responsive genes together, enriched biological functions were dominated by signaling- and membrane-associated processes, including phospholipid transport and translocation, protein kinase activity, voltage-gated calcium-channel complexes, transcriptional regulation, and DNA replication-associated processes (Figure 6C; Table S17).

**Figure 6:**
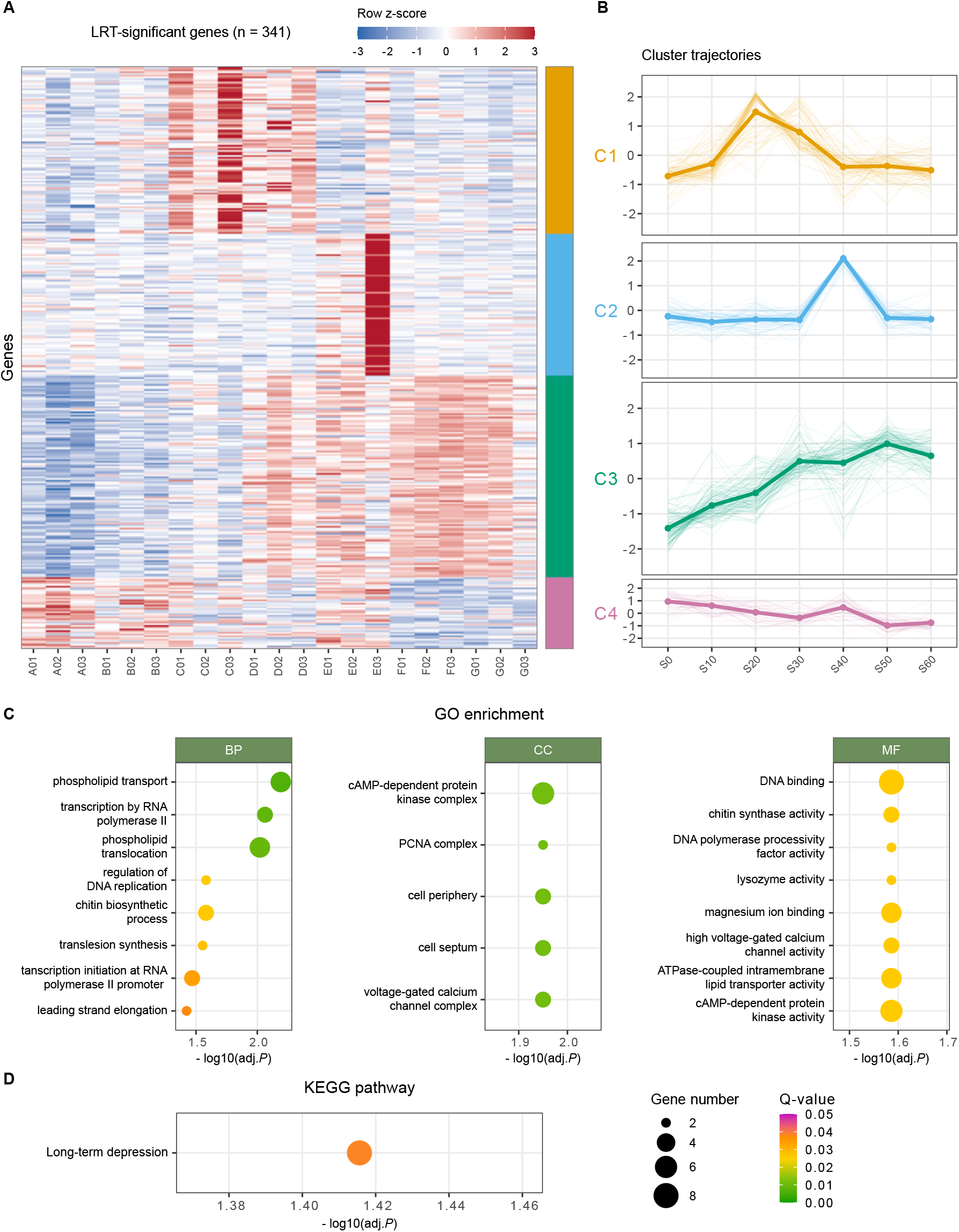
Transcriptional dynamics of habituation in *Stentor coeruleus*. **(A)** Heatmap of the 341 genes with significant expression change across the stimulus series (likelihood-ratio test, adjusted *p <* 0.05); rows are genes (row z-scored), columns are the 21 libraries grouped by time point, and the right-hand color bar marks four k-means clusters (C1–C4). **(B)** Mean expression trajectories of the four clusters across the stimulus series (S0–S60); thin lines, individual genes; thick lines, cluster means. **(C)** GO enrichment among the 341 time-course genes for biological process (BP), cellular component (CC), and molecular function (MF); dot size indicates the number of genes and dot color the adjusted *p*-value. **(D)** KEGG pathway over-representation; long-term depression was the only significantly enriched pathway. Enrichment was tested against the annotated *S. coeruleus* gene set as background; full results are provided in Table S17.

### 2.8 Habituation-responsive genes are enriched for a long-term-depression-associated signaling program

Given the prominent enrichment of signaling-related functions, we next asked whether habituation-responsive genes converged on specific signaling pathways. KEGG pathway enrichment analysis of the complete set of 341 habituation-responsive genes identified long-term depression (LTD), a signaling pathway classically associated with learning-related synaptic plasticity in animals ^25^, as the only significantly enriched KEGG pathway (Figure 6D).

Eight habituation-responsive genes were initially assigned to the LTD pathway. Following manual inspection of protein domain architectures, one anoctamin/TMEM16-family membrane protein was excluded because it is not a canonical component of the conserved LTD signaling module, leaving seven bona fide pathway components: five cGMP-dependent protein kinase (PKG1) paralogs, one voltage-gated calcium channel, and one protein phosphatase 2A (PP2A) regulatory subunit (Table S18; Figure S6). Domain architecture confirmed that each PKG paralog contained the characteristic fused cyclic nucleotide-binding and protein kinase domains expected of cGMP-dependent protein kinases ^26,27^. Notably, all seven curated pathway components belonged to the sustained activation cluster (C3) and showed progressively increasing expression throughout the stimulation series (Figure S6A). The coordinated up-regulation of these conserved pathway components defines a PKG–calcium channel–PP2A signaling module corresponding to the core components of the long-term depression pathway during habituation.

## 3 Discussion

Behavioral plasticity is often viewed through the lens of nervous systems, yet its underlying molecular principles may predate the evolution of neurons themselves ^28,29^. By integrating an improved reference genome with time-resolved transcriptomics, our study identifies a signaling architecture associated with habituation in the giant ciliate *Stentor coeruleus*. The prominent components, cyclic nucleotide-dependent kinases, a voltage-gated calcium channel, and a PP2A regulatory subunit, correspond to conserved intracellular elements represented in the neuronal long-term depression (LTD) pathway. Rather than implying a neuron-specific mechanism, this correspondence suggests that cyclic nucleotide signaling, calcium regulation, and reversible protein phosphorylation may constitute an ancient molecular framework for adaptive behavioral responses.

LTD emerged as the only significantly enriched KEGG pathway among habituation-responsive genes. Together with long-term potentiation (LTP), LTD comprises a broad class of activity-dependent processes that produce lasting decreases or increases in synaptic efficacy and contribute to learning and memory^25,30,31^. The direction of plasticity is determined not by a single universal pathway but by the magnitude and duration of Ca^2+^ signals, the balance of kinase and phosphatase activity, and the cellular targets through which these signals alter membrane-protein abundance, excitability, or transmitter release. NO–cGMP signaling, PKG, serine/threonine phosphatases, and voltage-gated calcium channels can therefore participate in either potentiation or depression, depending on cellular context and downstream substrates ^25,31–34^.

The LTD enrichment in *S. coeruleus* should therefore be interpreted at the level of shared signaling architecture rather than neuronal mechanism. The KEGG pathway is organized around a synaptic context, whereas the identities and functions of corresponding homologs in *Stentor* remain largely unresolved. Nevertheless, the conserved intracellular components shared between the two systems point to common molecular principles: repeated stimulation is accompanied by coordinated engagement of cyclic nucleotide signaling, calcium regulation, and reversible protein phosphorylation as responsiveness declines. The relevant substrates and the direction of regulation, however, cannot be inferred from transcript abundance alone.

Our findings also complement the recent study of Rajan et al. (2026), which independently implicated calcium-dependent phosphorylation in *S. coeruleus* habituation through pharmacological perturbation, transcriptomic, proteomic, and RNAi approaches. Although both studies investigate the same behavioral phenomenon, their experimental designs differ in several respects. Rajan et al. collected RNA from populations of stimulated cells across prolonged training and recovery paradigms, whereas we continuously tracked individual cells throughout a standardized habituation assay and selectively collected only those cells that remained substrate-attached and experienced the complete stimulation protocol up to each sampling point. This ensured that every sequenced cell had a defined stimulation history. Despite these methodological differences, both studies point to calcium-dependent signaling and reversible protein phosphorylation as prominent features of the habituation response.

The two studies differ in their emphasis on the kinases involved. Rajan et al. highlighted calcium-responsive protein kinases, whereas our pathway-level analysis identified sustained transcriptional activation of five PKG-like cyclic nucleotide-dependent kinases together with a voltage-gated calcium channel and a PP2A regulatory subunit. An interesting observation from genome annotation concerns the proteins discussed by Rajan et al. as CaMKII homologs. Their domain architectures are also compatible with CDPK/CDPK-like kinases: they contain C-terminal EF-hand Ca^2+^-binding motifs but lack a detectable CaMKII association domain required for canonical CaMKII holoenzyme assembly^35^ (Figure S6F). Consistent with this interpretation, neither the previous nor the current *S. coeruleus* annotation contains a protein with the complete canonical CaMKII architecture. Because KN-93 perturbs Ca^2+^/calmodulin-dependent signaling more broadly and does not uniquely diagnose canonical CaMKII activity^36^, these observations do not alter the central conclusion of Rajan et al. that calcium-dependent phosphorylation contributes to habituation. They instead suggest that the relevant calcium-responsive kinases in *S. coeruleus* may belong to the CDPK lineage. More broadly, the two studies converge on regulated kinase–phosphatase activity, rather than a single kinase family, as a central feature of habituation.

The cyclic nucleotide-dependent kinases identified here are particularly compelling because PKG is one of the most evolutionarily conserved regulators of behavioral plasticity. PKG orthologs occur throughout eukaryotes, from protists to mammals, while retaining remarkably conserved regulatory and catalytic architectures ^27,37^. Across animals, PKG has repeatedly been implicated in experience-dependent behavioral modification. The *Caenorhabditis elegans* ortholog EGL-4 regulates olfactory adaptation ^38^; the *Drosophila* foraging gene controls habituation of the proboscis-extension and giant-fiber escape responses and influences food-related behavioral plasticity^39–42^; and the honeybee ortholog *Amfor* contributes to the transition to foraging behavior ^43^. Although these behaviors differ substantially, they collectively illustrate a recurring evolutionary role for PKG in tuning behavioral responses according to previous experience or internal physiological state.

The classification of the *Stentor* proteins nevertheless requires some caution. PKA and PKG catalytic domains are closely related within the AGC kinase group and cannot be distinguished reliably from the kinase domain alone ^34,44^. The proteins identified here contain tandem N-terminal cyclic nucleotide-binding domains fused to a C-terminal AGC kinase domain (Figure S6B), an arrangement characteristic of PKG, whereas canonical PKA comprises separate regulatory and catalytic subunits. This fused architecture, together with sequence similarity and pathway annotation, supports their description as PKG-like kinases. Direct biochemical measurements of nucleotide selectivity and kinase activation will nevertheless be required to establish cGMP dependence and exclude divergent PKA-related signaling.

The connection to PKG is especially relevant because cGMP already has a documented behavioral role in *S. coeruleus*. Pharmacologically elevating intracellular cGMP or inhibiting its degradation reduces photophobic responsiveness and prolongs response latency, whereas manipulations expected to lower cGMP produce the opposite effect^45^. Those experiments established that cGMP can down-regulate a sensory response in *Stentor* but did not identify its downstream effector. The coordinated up-regulation of five PKG-like paralogs during mechanical habituation now nominates this branch of cyclic nucleotide signaling as a candidate contributor to the progressive decline in responsiveness.

How this module influences contraction remains unresolved. Rapid contraction in *S. coeruleus* is driven by Ca^2+^-responsive myonemes containing centrin-family proteins ^46,47^. Increased expression of a voltage-gated calcium channel might therefore appear inconsistent with habituation, because greater channel activity would be expected to promote Ca^2+^ influx and contraction. In vascular smooth muscle, however, PKG reduces contractility by limiting voltage-gated Ca^2+^ entry, suppressing IP_3_-dependent Ca^2+^ release, promoting Ca^2+^ sequestration, and lowering the Ca^2+^ sensitivity of the contractile machinery^27,34^. Increased channel transcription in *Stentor* could therefore represent a homeostatic response to reduced channel activity or enhanced Ca^2+^ clearance under elevated PKG signaling. This interpretation remains speculative, because transcript abundance does not establish channel activity, kinase activity, or net Ca^2+^ flux. Direct Ca^2+^ imaging, measurements of channel function, selective perturbation of the candidate kinases, and identification of their phosphorylation substrates in *Stentor* will be needed to determine whether this module dampens or enhances excitability during habituation.

The anoctamin/TMEM16-family protein excluded from the curated LTD set remains independently interesting. TMEM16 proteins include Ca^2+^-activated ion channels, phospholipid scram-blases, and dual-function proteins. Canonical ANO1/TMEM16A and ANO2/TMEM16B function as Ca^2+^-activated chloride channels, whereas other family members primarily reorganize membrane phospholipids ^48–50^. The *Stentor* protein therefore cannot be assigned as a chloride channel from homology alone. It may instead contribute to calcium-dependent regulation of cortical membrane organization, trafficking, surface charge, or excitability; direct assays of ion transport and phospholipid scrambling will be required to establish its molecular function.

Pathway-level interpretation also has important limits. Enrichment identifies statistical over-representation of annotated components, not activation of a complete pathway or functional requirement for its members. Moreover, only approximately 47% and 39% of predicted genes carry GO and KEGG annotations, respectively, leaving unannotated and *S. coeruleus*-specific factors invisible to these analyses. The likelihood-ratio test may also miss genes altered only at individual time points and cannot capture regulation through translation, localization, phosphorylation, or other post-transcriptional mechanisms. The present analysis therefore defines a focused set of candidate regulators rather than a complete mechanistic account of habituation.

Beyond the molecular insights into habituation, this study also establishes a substantially improved genomic framework for *S. coeruleus*. The previously available genome, assembled from Illumina short reads ^11^, was necessarily constrained by the sequencing technologies available at the time, limiting assembly contiguity and analyses of genome architecture. By leveraging PacBio HiFi sequencing together with modern long-read assembly approaches, SteCoeF v1 substantially improves genome continuity and annotation quality, enabling robust reconstruction of intragenomic collinearity, resolution of two rounds of whole-genome duplication, and more reliable inference of lineage-specific gene-family evolution. Although these evolutionary analyses do not establish a causal relationship between genome evolution and habituation, they provide an important framework for future studies investigating how duplicated genes have diversified and whether they contribute to behavioral plasticity in *Stentor*. More broadly, the improved reference genome provides a valuable resource for comparative and functional genomics across ciliates.

Taken together, our study establishes an improved reference genome for *S. coeruleus* and links this resource to the molecular analysis of a classical form of single-cell learning. By integrating genome assembly, comparative genomics, and time-resolved transcriptomics, we identify a conserved cyclic nucleotide-dependent signaling module associated with habituation while providing an evolutionary framework for interpreting its origins. More broadly, these findings establish *S. coeruleus* as a uniquely tractable model for investigating how conserved signaling networks enable adaptive behavior in the absence of neurons, providing a window into the molecular foundations from which learning-related mechanisms may have evolved.

## 4 Methods

### 4.1 *Stentor* culturing

*Stentor coeruleus* culturing followed procedures described previously^51^. Briefly, *S. coeruleus* were obtained from Carolina Biological (catalog no. 131598) and maintained in filtered spring water (Carolina Biological, catalog no. 132450) for a minimum of two weeks before experimentation. Cultures were kept in Pyrex bowls in dark drawers at 22°C when not in use. *Chlamydomonas reinhardtii* (Chlamydomonas Resource Center, CC-125) were grown on 1.5% agar plates prepared with TAP medium (PhytoTech Labs, T8224) under continuous bright light. For feeding, Chlamydomonas cells were transferred from agar plates to 50 mL of filtered spring water using five loopfuls of a sterile inoculating loop, and large clumps were dispersed by gentle pipetting and shaking. *Stentor* cultures (500 mL) were fed with 7 mL of this feeding solution every four to five days. *Stentor* and *Chlamydomonas* cultures were inspected weekly for contamination, cell death, or reduced growth, and *Stentor* cultures were transferred to fresh spring water monthly to maintain culture health and minimize bacterial overgrowth.

### 4.2 Genomic DNA extraction and PacBio HiFi sequencing

Cultured *Stentor coeruleus* cells were collected and washed repeatedly in filtered spring water to remove food organisms and debris, then concentrated by centrifugation at 1,000 rpm for 5 min. The supernatant was removed, leaving approximately 100 µL of packed cells. Genomic DNA extraction, sample quality control, library preparation, and PacBio Revio HiFi sequencing were performed by the Bauer Core Facility (Harvard University). High-molecular-weight DNA was extracted with the NanoBind PanDNA kit (Pacific Biosciences) following the HMW DNA extraction protocol for cultured suspension cells, eluting in 100 µL of LTE buffer; DNA was quantified with the Qubit dsDNA HS assay (Thermo Fisher Scientific) and sized on a 165-kb Femto Pulse assay (Agilent). Fragments below 10 kb were depleted with Short Read Eliminator (Pacific Biosciences; 1-h extended centrifugation), and the DNA was sheared in 200 µL at speed 30 on a Megaruptor 3 (Diagenode). SMRTbell libraries were constructed from the sheared DNA with the SMRTbell prep kit 3.0 (Pacific Biosciences) using the whole-genome and metagenome protocol, including enzymatic DNA-damage repair and A-tailing, blunt-adapter ligation, nuclease treatment to remove unligated fragments and adapters, and AMPure PB bead cleanup. The library was then prepared with the Revio SPRQ polymerase kit and standard sequencing primer (annealing–binding–cleanup workflow) and sequenced on a 25M Revio SMRTcell (280 pM on-plate loading concentration; 30-h movie) on the PacBio Revio system, generating HiFi (circular consensus) reads. Sequencing yielded 9,125,848 HiFi reads (98.67 Gb; mean read length 10.8 kb, read N50 12.6 kb, median read quality Q40).

### 4.3 Genome assembly

To evaluate the effect of assembler choice, three HiFi-compatible assemblers were compared: Flye v2.9.6^13^, hifiasm v0.25.0^15^, and HiCanu ^14^. Flye was run with `--genome-size 80m` and `–threads 16` (and `--asm-coverage 100` for the full, high-coverage dataset), in `--pacbio-hifi` modes tested with `--read-error 0.05`. hifiasm was run in primary mode (`-o <prefix> -t <threads> --primary`), with `--hom-cov 1000` on the full dataset, and primary contigs were extracted from the `p_ctg` graph. HiCanu (Canu v2.3) was run with `genomeSize=80m`, `-pacbio-hifi`, homopolymer compression (`-homopolycompress`), and a 1-kb minimum read length. The final reference assembly was produced with Flye v2.9.6 `--pacbio-hifi` on the full HiFi dataset, which yielded a compact assembly.

### 4.4 Assembly curation and decontamination

The primary Flye assembly was curated in the order purge dups, Inspector, then contamination screening. Haplotypic and duplicate contigs were removed with purge dups v1.2.5^52^; following the purge dups documentation, the default cutoff was applied first, followed by manually adjusted cutoffs derived from the read-depth histogram (for high-coverage data, pbcstat was run with -M 10001, and cutoffs were set manually because calcuts v1.2.5 segfaulted on this dataset). Base-level errors and structural misassemblies were then corrected with Inspector v1.3^53^. Finally, adapters, vectors, and foreign or endosymbiotic sequences were screened with NCBI’s Foreign Contamination Screening tools FCS-adapter and FCS-GX^54^, each run with default settings and the taxonomic identifier set to that of *S. coeruleus*. Because ribosomal DNA (rDNA) contigs also carry protein-coding genes, contamination and rRNA handling used position-based filtering rather than whole-contig exclusion. After screening, the curated assembly was re-evaluated by mapping the raw HiFi reads back to it with Inspector, which reported a final consensus quality value (QV) of 54.38 (improved from 47.82 for the pre-curation assembly) and a single residual structural error (reduced from 981). The final assembly, SteCoeF v1, comprised 185 contigs spanning 60.18 Mb (contig N50 445 kb, N75 340 kb, longest contig 949 kb).

### 4.5 Independent iterative assembly and telomere identification

As an orthogonal assembly for cross-validation, a telomere-to-telomere-oriented assembly (Ste-CoeH v1) was generated following the strategy of Wang et al. (2021). Twenty random 60× subsets of the HiFi reads were each assembled with hifiasm in primary mode (`-o <prefix> -t <threads> --primary`), and primary contigs were extracted from the `p ctg` graph. Telomeres were identified using the heterotrich motifs 5*^→^*-CCCTAACA and 3*^→^*-TGTTAGGG; a contig end was scored as telomere-capped when at least 10 consecutive repeat units occurred within 1,000 bp of the terminus, and contigs capped at both ends were classified as closed. Beginning from one subset assembly, closed contigs were retained while the remaining unclosed contigs were merged iteratively, round by round, against the other subset assemblies with quickmerge (`merge wrapper.py -l 130000 -ml 10000`; i.e., a 130-kb anchor-contig length cutoff and a 10-kb minimum alignment length), accumulating closed contigs across rounds. The merged assembly was then curated with custom scripts to remove misjoins and redundant contigs. SteCoeH v1 (382 contigs, 86.02 Mb, contig N50 378 kb, with 59.2% of contigs telomere-capped at both ends) served only as an independent reference for cross-validation: SteCoeF v1 and SteCoeH v1 share 99.62% sequence identity, with ~90.1% of SteCoeF v1 bases represented in SteCoeH v1 and 99.1% of SteCoeH v1 aligning to SteCoeF v1.

### 4.6 Assembly evaluation and comparison

All assemblies were evaluated for contiguity (number of contigs, assembly size, N50, longest contig) and completeness. Completeness was assessed with BUSCO v5.4.6 against the alveolata odb10 lineage ^55^ (*n* = 171). Contiguity and correctness metrics were summarized with QUAST v5.2.0^56^, and per-base accuracy (QV) and structural errors were quantified by mapping the HiFi reads back to each assembly with Inspector ^53^. High duplicated-BUSCO content was interpreted as biological signal from WGD rather than as assembly error. Pairwise assembly-to-assembly comparisons (SteCoeF v1 versus SteCoeH v1 and versus the 2017 assembly) were performed with nucmer and dnadiff from MUMmer v4^57^ to obtain identity and one-to-one alignment coordinates, and large-and small-scale structural differences were quantified with NucDiff v2.0.3^58^.

### 4.7 Repeat annotation

Repetitive elements were modeled de novo with RepeatModeler v2.0.8 (search engine rmblast 2.17.1+) ^59^ on SteCoeF v1, which yielded 112 repeat families; structural analysis detected no LTR retrotransposons. The genome was then screened against this custom library with RepeatMasker v4.2.3^60^, which masked 7.28% of the assembly: 5.99% interspersed repeats, predominantly unclassified elements (5.25%), with retroelements (0.57%) and DNA transposons (0.18%) each contributing little, together with 0.86% simple repeats and 0.18% low-complexity sequence. This low repeat content is consistent with that of the 2017 assembly^11^. No soft-masking was applied prior to gene prediction, and structural annotation proceeded on the unmasked assembly.

### 4.8 Structural annotation

RNA-seq reads were aligned to support gene prediction with HISAT2^61^, which was recompiled from source to accommodate the exceptionally short introns of *Stentor* : the minimum intron length in`hisat2.cpp` was reduced from 20 to 9, and the recompiled binary was run after loading `gcc/12.2.0-fasrc01`. Alignments used `--max-intronlen 101` and `--very-sensitive` and were sorted and indexed with SAMtools v1.21^62^. Protein-coding genes were predicted with Intronarrator ^18^, which detects and removes introns, runs AUGUSTUS v3.5.0^63^ with an intronless model, and reinserts introns; Intronarrator was run with `MAX INTRON LEN = 101` and `GENETIC CODE = 1` (the standard code, as *Stentor* does not use a ciliate-specific code). Contigs and genes were renamed to the SteCoeF prefix using a modified GFF parser and custom scripts. Transfer RNAs were annotated with tRNAscan-SE v2.0.12^64^, and ribosomal RNA loci were located with cmsearch (Infernal v1.1.5) ^65^ against Rfam for use in position-based read filtering (below). Structural annotation yielded 32,306 protein-coding genes.

### 4.9 Functional annotation

Predicted proteins were functionally annotated by combining four complementary tools, applying a default e-value cutoff of `1e-6` unless otherwise stated. Orthology and preliminary Gene Ontology (GO) and KEGG Orthology (KO) assignments were obtained with eggNOG-mapper v2.1.13 (`--itype proteins --tax scope Eukaryota --go evidence non-electronic`) ^66^. Protein domains, families, and GO terms were assigned with InterProScan v5.70-102.0 (applications `Pfam`, `PANTHER`, `TIGRFAM`, `CDD`, `SFLD`, `ProSiteProfiles`, and `Hamap`, with `-goterms -pa`) ^67^. Homology-based descriptions were obtained with DIAMOND v2.1.12 (`--more-sensitive --max-target-seqs 1`, e-value `1e-5`) ^68^ against SwissProt, TrEMBL, and NCBI NR (downloaded April 15, 2023), giving priority to SwissProt/TrEMBL matches above 95% identity. KO terms were assigned with KofamScan v1.3.0 (significant hits only) ^69^ with an e-value cutoff of `1e-9`. The four sources were integrated per gene with explicit precedence rules: descriptions were taken in the order SwissProt (high-identity) *>* eggNOG *>* InterPro *>* “hypothetical protein”. To avoid the annotation inflation that arises when eggNOG propagates the annotation union of an entire orthologous group, GO terms used for all downstream analyses were taken from InterProScan only, and KO terms were assigned with KofamScan taking precedence, with eggNOG used only to fill genes lacking a KofamScan KO. Across the genome, 93.3% of genes received some annotation; GO terms (InterProScan-only) were assigned to 15,295 genes (47.3%; mean 3.3 GO terms per annotated gene), and KO terms to 12,513 genes (38.7%).

### 4.10 Whole-genome duplication analysis

Intragenomic collinearity was analyzed to test for WGD. All-versus-all protein alignments (BLASTP) were supplied to MCScanX^19^, retaining collinear blocks containing more than five genes. Synonymous-substitution rates (*K_s_*) between syntenic paralog pairs and the maximum collinear-duplication depth were computed with WGDI v0.6.4^70^. Across 480 collinear blocks, 7,125 syntenic paralog pairs (11,158 genes) were identified. Two lines of evidence supported two successive rounds of WGD: a bimodal Ks distribution with peaks at ~1.70 and ~4.01, and a maximum collinear-duplication depth of three. Collinearity was visualized as a Circos plot ^71^, with contig order optimized so that collinear contigs were grouped within the same sector.

### 4.11 Gene-family evolution

Orthologous groups were inferred across a panel of ciliates with OrthoFinder v2.5.5^72^. The input proteomes comprised *Stentor coeruleus* (this study), *Stentor roeselii* (obtained from the authors of Zheng et al., 2025), *Stentor pyriformis* (StentorDB,https://stentor.ciliate.or `g`), *Oxytricha trifallax* (GCA 000295675.1), *Stylonychia lemnae* (GCA 000751175.1), *Paramecium tetraurelia* (GCA 000165425.1), *Tetrahymena thermophila* (GCA 000189635.1), and *Pseudocohnilembus persalinus* (GCA 001447515.1). Of 195,462 input proteins, 170,034 (87.0%) were assigned to 31,185 orthogroups. An ultrametric species tree was generated from single-copy orthologs, and divergence times were estimated with r8s ^23^, calibrated using node ages from the TimeTree database ^73^. Gene-family expansions and contractions were then modeled along this tree with CAFE5^22^. Expanded and contracted families were tested for GO and KEGG enrichment with clusterProfiler ^74^. Because gene-family counts are sensitive to assembly contiguity and annotation completeness, directional expansion/contraction contrasts were interpreted only within this pipeline and not compared in absolute terms across studies.

### 4.12 Habituation behavioral assay

Habituation was assayed in individually anchored *Stentor coeruleus*. Cells attached to the substrate were subjected to automatically delivered mechanical taps at a frequency of one tap per minute, using the delivery apparatus and imaging setup described for Figure 5. The contractile response of each cell to each tap was scored from video recordings. For behavioral analysis, cells were tracked across the stimulus series, and cells that detached or moved before completion of the analyzed series were excluded. For each replicate, the fraction of eligible cells contracting was quantified at each stimulus number (*n* = 36–114 cells per replicate), yielding the behavioral habituation curve.

### 4.13 RNA sample preparation and sequencing

Twenty-one RNA-seq libraries were generated, corresponding to seven points along the stimulus series (S0, S10, S20, S30, S40, S50, and S60) with three biological replicates each; each replicate was an independent batch of pooled cells. Total RNA was extracted with the PureLink RNA Mini Kit (Thermo Fisher Scientific, 12183018A), treated with ezDNase enzyme (Invitrogen, 11766051) to remove residual genomic DNA, and purified with RNAClean XP beads (Beckman Coulter, A66514). Each sample was collected from the cells assayed in the corresponding behavioral replicate, so that the number of cells pooled per sample matches the per-replicate n shown in Figure 5E 36–114 cells per sample). Library preparation and sequencing were performed by the Bauer Core Facility (Harvard University). RNA integrity was assessed on a TapeStation 4200 (Agilent), RNA was quantified with the RiboGreen assay (Thermo Fisher Scientific), and samples were normalized to 20 ng. Polyadenylated mRNA was captured on oligo-dT magnetic beads, fragmented to 300-400 bp, and converted into strand-specific libraries by dUTP incorporation in the second strand using the Watchmaker mRNA library prep kit (Watchmaker Genomics) with xGen stubby adapters (Integrated DNA Technologies). Libraries were amplified for 12 cycles with Equinox master mix (Watchmaker Genomics) and dual 8-bp indices (Integrated DNA Technologies); all volumes were miniaturized to one-fifth reactions on a Mantis liquid handler (Formulatrix), and bead-based cleanups were performed on a SciClone G3 NGSx workstation (PerkinElmer). Final libraries were assessed on a TapeStation 4200 (Agilent), quantified by qPCR (Roche), pooled, and sequenced on half a lane of a NovaSeq X Plus 10B flow cell with paired-end 150-bp reads (Illumina).

### 4.14 RNA-seq read processing and quantification

Reads from the 21 libraries (7 time points × 3 replicates) were aligned to SteCoeF v1 with the recompiled HISAT2^61^ (`--max-intronlen 101 --very-sensitive --no-unal`), yielding a mean alignment rate of 97.4%. Because the macronuclear genome carries rDNA loci that also contain protein-coding genes, ribosomal reads were removed by position: rDNA coordinates identified by cmsearch were used to filter overlapping reads with BEDTools ^75^, rather than excluding entire rDNA-bearing contigs. Reads were counted per gene with featureCounts ^76^ using `-p` --countReadPairs -s 2 -t gene -g ID --primary, giving a mean of 93.7% assigned read pairs. Genes with at least 10 counts in at least 3 samples were retained for downstream analysis.

### 4.15 Differential expression and temporal clustering

Differential expression was analyzed with DESeq2 v1.46.0^77^. For quality control and visualization, counts were transformed with the variance-stabilizing transformation (VST). To identify genes that changed across the full stimulus series rather than at a single contrast, a likelihood-ratio test (LRT) was used, comparing the full model (`∼condition`) with a reduced model (`∼1`); genes with adjusted *p<* 0.05 (Benjamini–Hochberg) were considered significant, with no fold-change threshold (the LRT tests overall variation rather than direction). This identified 341 habituation-responsive genes. These genes were grouped by temporal trajectory using k-means clustering (*k* = 4, *nstart* = 50, `set.seed(42)`) on *z*-scored, per-condition VST means, yielding four interpretable programs: a transient cluster peaking at S20 (*n* = 98), a transient cluster peaking at S40 (*n* = 83), a sustained, progressively up-regulated cluster (*n* = 118), and a slowly down-regulated cluster (*n* = 42).

### 4.16 Functional enrichment

Over-representation analysis of the LRT-significant genes and of each temporal cluster was performed with clusterProfiler ^74^. Because *Stentor* is not represented in OrgDb or as a KEGG organism, enrichment used the enricher() function with custom TERM2GENE tables built from the InterProScan GO assignments and the KO assignments; GO term names were retrieved from GO.db v3.20.0. To avoid spuriously inflating the background, the gene universe for each test was the set of genes tested by DESeq2 intersected with the set of annotated genes, rather than all genes in the genome. *p*-values were adjusted by the Benjamini–Hochberg method (significance at adjusted *p <* 0.05). The only enriched KEGG pathway among the habituation-responsive genes was long-term depression (map04730; adjusted *p* = 0.038).

Several annotation caveats were handled explicitly. KO terms were assigned with KofamScan taking precedence; for genes whose only KO came from eggNOG, multi-KO assignments (which arise when eggNOG propagates the annotation union of an orthologous group) were reconciled against Pfam/InterPro domain architecture and SwissProt best hits. The InterPro2GO mapping of the shared AGC-kinase catalytic domain assigns “cAMP-dependent protein kinase activity” (GO:0004691) without discriminating cGMP-from cAMP-dependent kinases; cGMP-dependent protein kinase (PKG) identity was therefore established from orthology (KEGG K07376) and domain architecture rather than from GO. For the long-term depression pathway, the eight genes contributing to the enrichment were curated at the domain level: five were confirmed as PKG1 paralogs (fused cyclic-nucleotide-binding and protein-kinase domains, best matches to characterized PKGs), one as a voltage-gated calcium channel, and one as a PP2A regulatory subunit, whereas an eighth gene assigned K07376 only through an eggNOG multi-KO annotation, but bearing anoctamin/calcium-activated chloride-channel domains, was reassigned and treated as a non-member. Over-representation statistics were not recomputed after this curation and correspond to the original KO assignments.

### 4.17 Data visualization and statistical analysis

Unless otherwise stated, statistical analyses and figures were produced in R v4.4.2 using DESeq2, clusterProfiler, ggplot2, and patchwork, and phylogenetic trees were drawn with ggtree. Figures were assembled and edited in Adobe Illustrator. Specific statistical tests, sample sizes, and significance thresholds are reported with each analysis above and in the corresponding figure legends.

### 4.18 Data and code availability

All raw sequencing data generated in this study have been deposited in the NCBI Sequence Read Archive under BioProject PRJNA1489669. The PacBio Revio HiFi whole-genome reads are available under run accession SRR39452315 (BioSample SAMN61407562), and the 21 poly(A) mRNA-seq libraries under run accessions SRR39453085–SRR39453105 (BioSamples SAMN61407585–SAMN61407605). All original code used for genome assembly, comparative genomics, and transcriptomic analyses is available at GitHub (https://github.com/yuemiao1017/Stentor_genome_habituation_transcriptomics). Code and data used for the protein domain reassessment of *Stentor* calcium-dependent kinases (CDPK/CDPK-like versus CaMKII) are available at GitHub (https://github.com/austenmtheroux/stentor_cdpk_vs_camkii_reassessment).

## 5 Acknowledgments

We thank Ashley R. Albright and Wallace F. Marshall for valuable discussions on *Stentor* biology. We are grateful to Tim Sackton for advice on genome assembly and annotation, Jie Xiong for guidance and helpful discussions on the iterative genome assembly strategy, and Yan Ying for generously sharing the *Stentor roeselii* genome assembly resources generated by her laboratory. We thank Claire Bailey Hartmann, Nicole El-Ali, Kelly Cribari and the Bauer Core Facility for assistance with sequencing and technical support. We also thank John Vastola, Zachary Kelso, and Madeleine Snyder for helpful discussions throughout this project.

This work was supported by a Polymath Award from Schmidt Sciences and the Air Force Office of Scientific Research (FA9550-22-1-0345). Part of this work was completed while SJG was in residence at the Kavli Institute for Theoretical Physics, supported in part by grant NSF PHY-2309135 and Gordon and Betty Moore Foundation Grant No. 2919.02.

## 6 Supplemental figures

**Figure S1:**
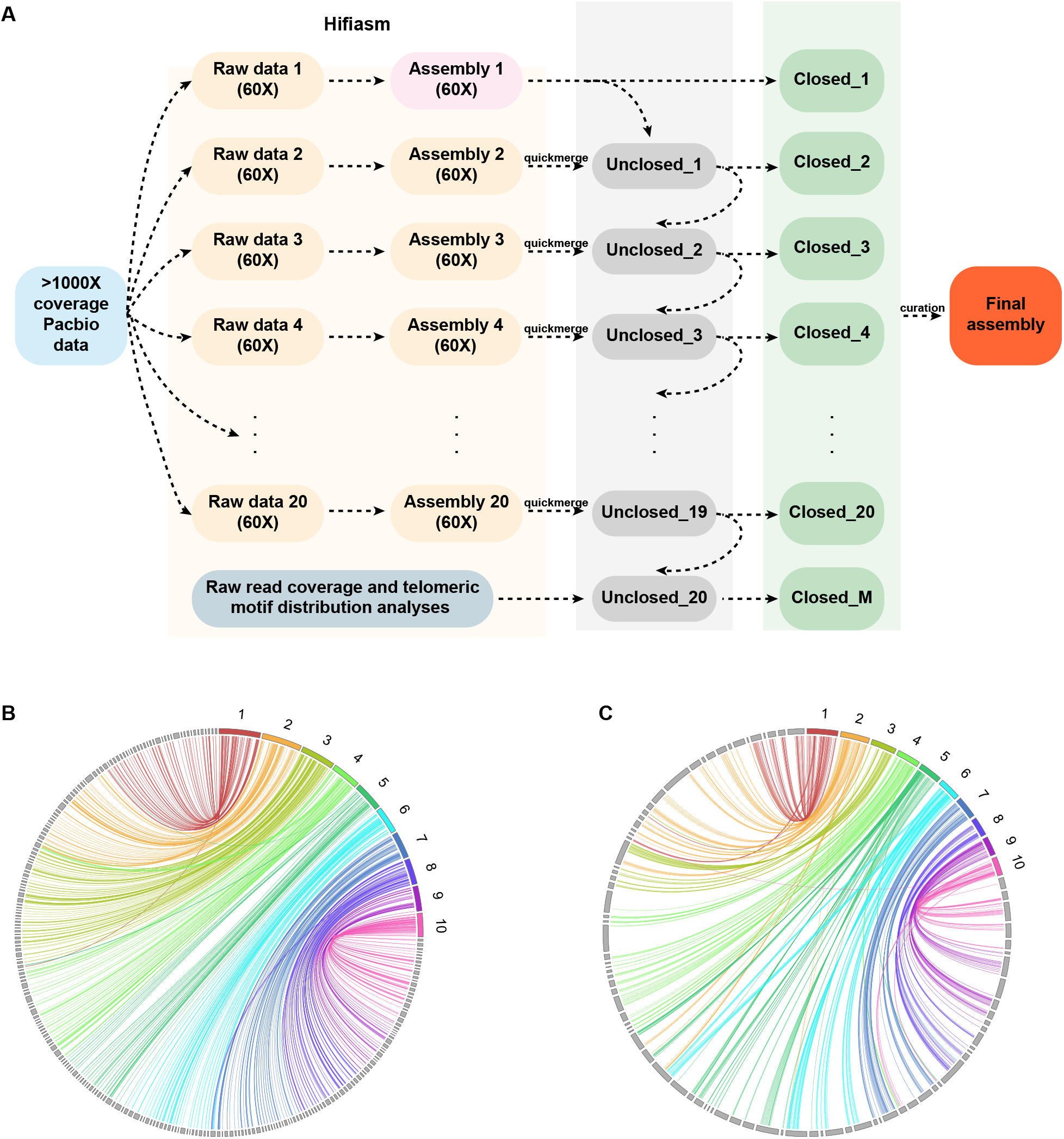
Iterative MAC genome assembly and cross-assembly validation. **(A)** Schematic of the iterative assembly strategy (adapted from Wang et al., 2021). High-coverage (*>*1,000×) PacBio data were partitioned into independent ~60× subsets, each assembled with hifiasm. In each round (1–20), contigs with *Stentor* -specific telomere motifs at both ends were classified “Closed”; contigs lacking terminal telomeres (“Unclosed 1–20”) were iteratively merged with quickmerge, followed by raw-read coverage and telomere-motif analyses and final curated steps. **(B)**Whole-genome alignment between SteCoeH v1 and the previously published 2017 Illumina assembly. **(C)** Whole-genome alignment between SteCoeF v1 and SteCoeH v1. In (B) and (C), colored contigs represent the reference assembly and grey contigs represent the comparison assembly. Links indicate aligned genomic regions and are colored according to the corresponding reference contig. Contigs are ordered by length, and only the ten longest reference contigs are labeled.

**Figure S2:**
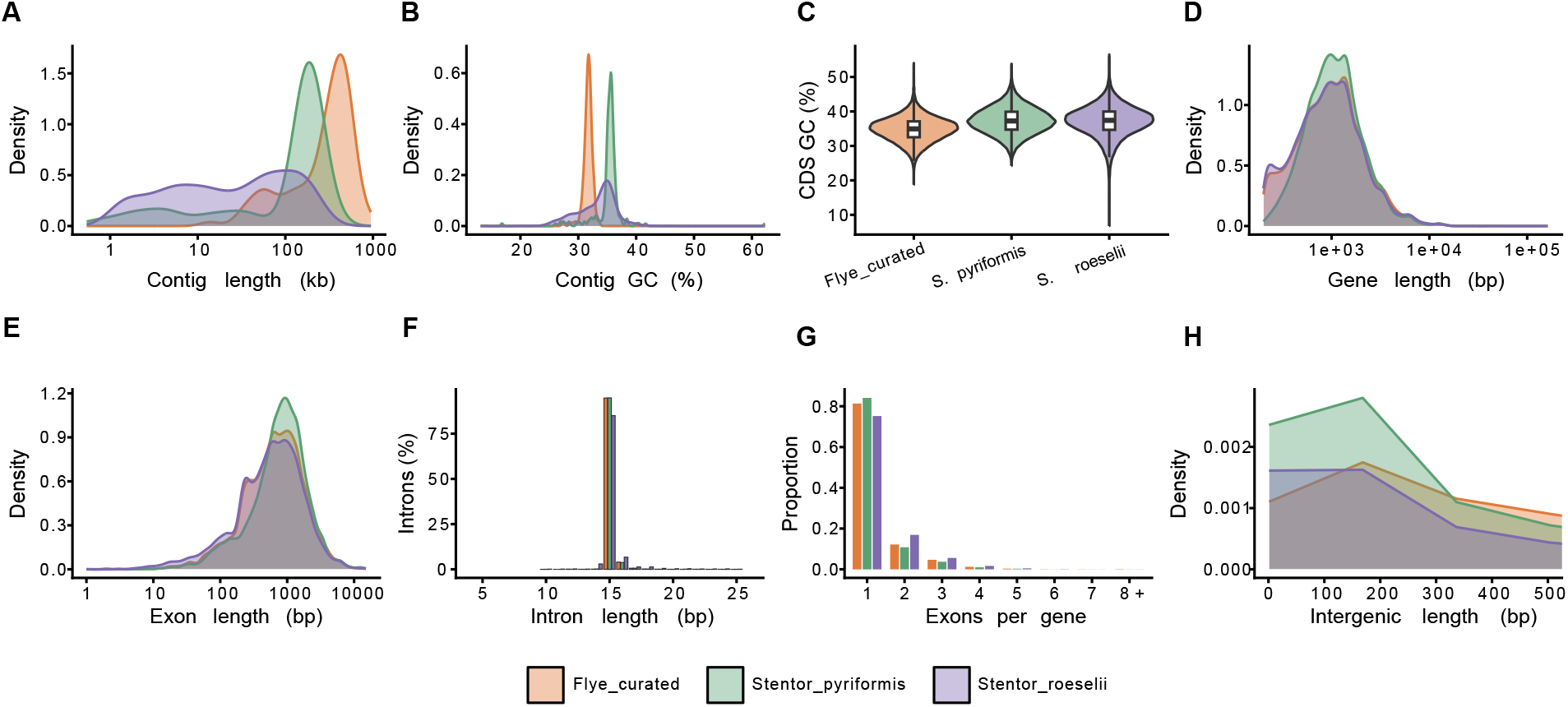
Comparison of genome and gene-structure features across three *Stentor* species. Genome- and gene-level features of *S. coeruleus* (SteCoeF v1; orange), *S. pyriformis* (green), and *S. roeselii* (purple): **(A)** contig length, **(B)** contig GC content, **(C)** CDS GC content, **(D)** gene length, **(E)** exon length, **(F)** intron length, **(G)** number of exons per gene, and **(H)** intergenic length. Annotations for *S. pyriformis* and *S. roeselii* were obtained from Boudreau et al. (2025) and Zheng et al. (2025), respectively. Numeric statistics for all three species are provided in Table S6.

**Figure S3:**
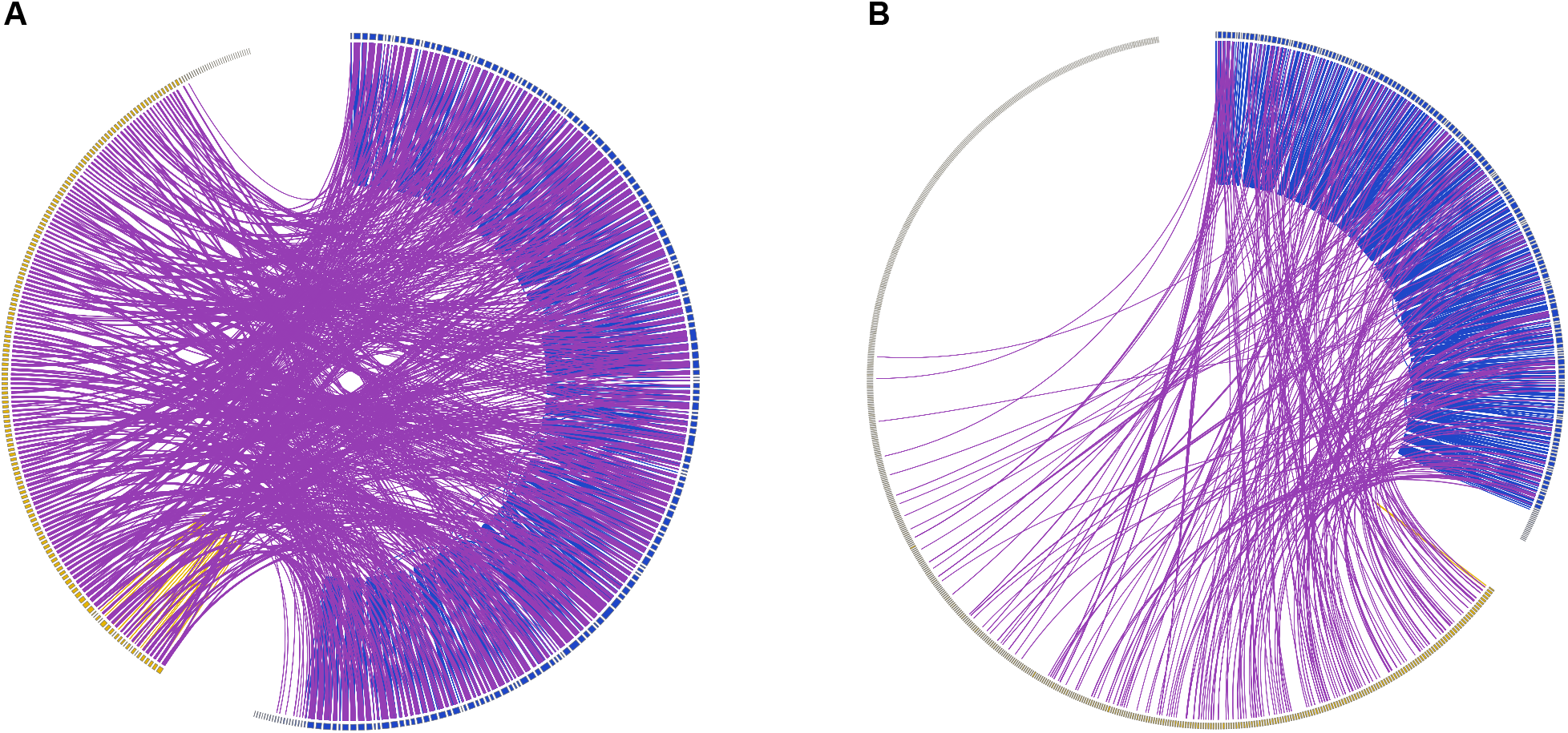
Intergenomic collinearity between *Stentor coeruleus* and two congeneric species. Circos plots showing between-genome collinearity for **(A)** *S. coeruleus* versus *S. pyriformis* and **(B)** *S. coeruleus* versus *S. roeselii*. In each plot, the two genomes are shown on opposing arcs separated by a gap. Links are colored by relationship type: between-species collinear pairs are shown in purple, and within-species collinear pairs are shown in blue or yellow.

**Figure S4:**
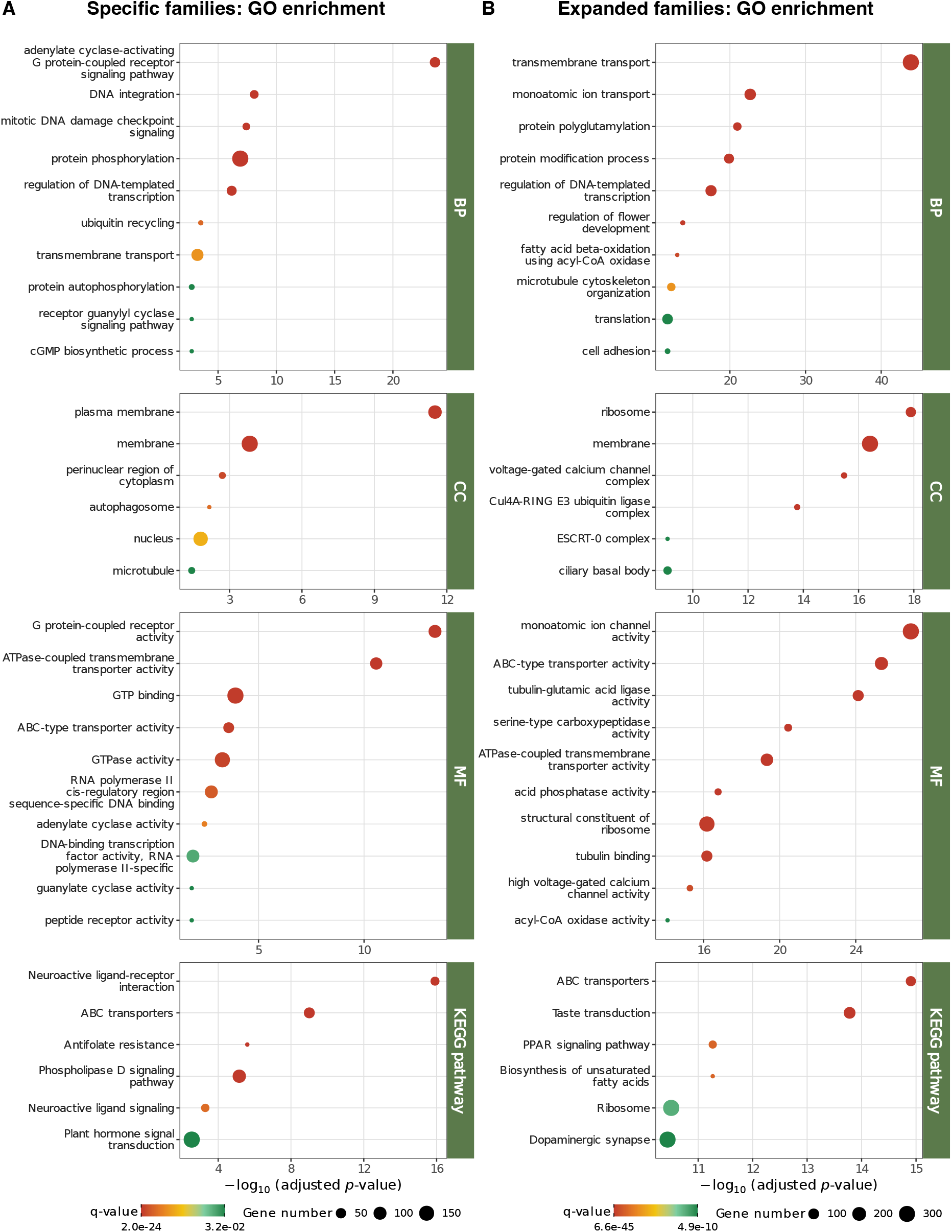
Functional enrichment of *Stentor coeruleus*-specific and expanded gene families. GO (biological process, BP; cellular component, CC; molecular function, MF) and KEGG pathway over-representation for **(A)** genes from *S. coeruleus*-specific orthogroups (5,232 genes; Table S12) and **(B)** genes from gene families significantly expanded on the *S. coeruleus* branch (2,922 genes; Table S13). Dot size indicates the number of genes associated with each term, dot color indicates the adjusted *p*-value (*q*), and the x-axis shows − log_10_(adjusted *p* value). Enrichment was tested against the annotated *S. coeruleus* gene set as background.

**Figure S5:**
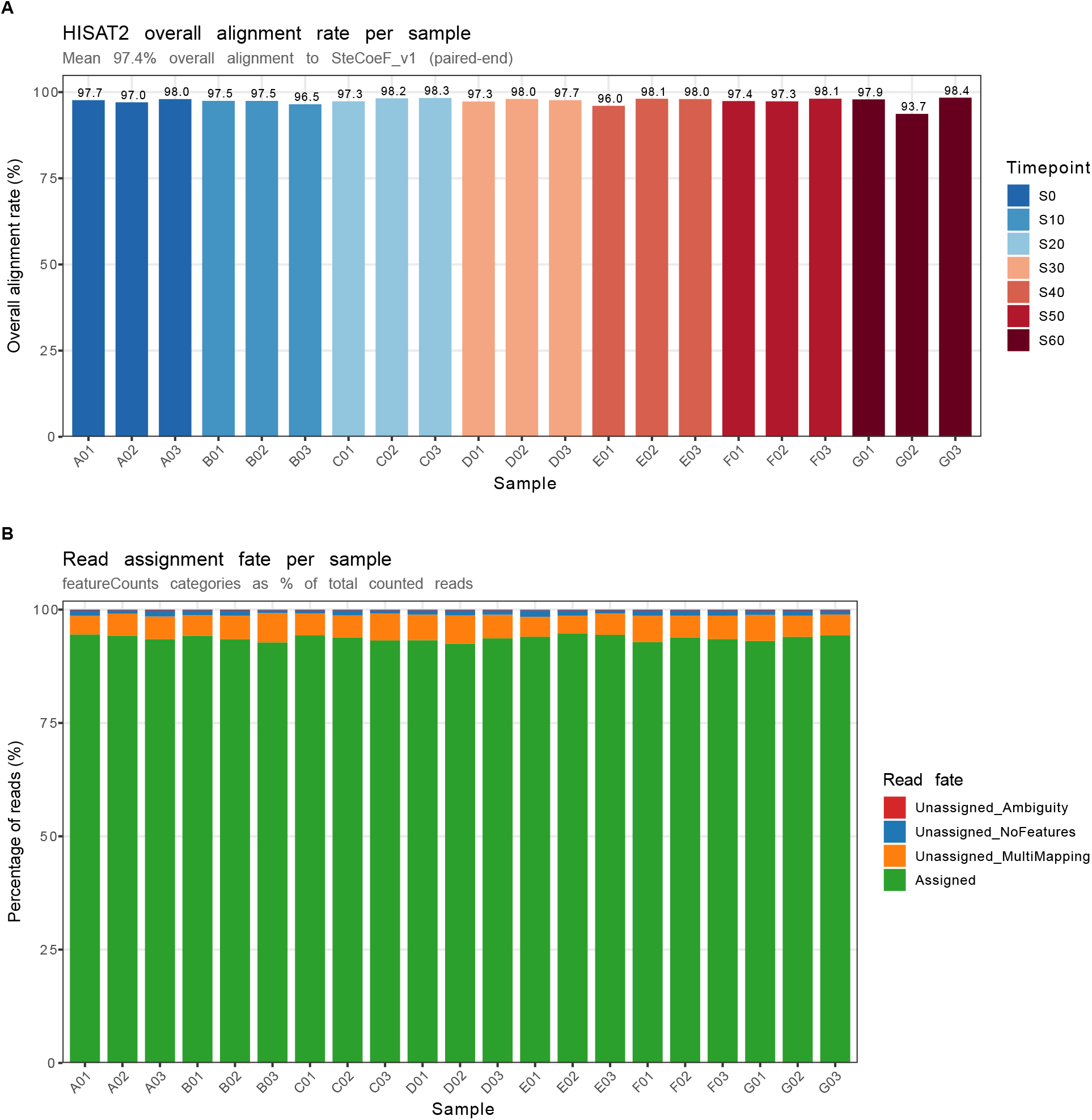
RNA-seq alignment and read-assignment quality control. **(A)** Overall HISAT2 alignment rate to the SteCoeF v1 reference for each of the 21 libraries (mean 97.4%; paired-end), colored by time point. **(B)** featureCounts read-assignment categories (assigned; unassigned owing to multimapping, no features, or ambiguity) as a percentage of counted reads per library. Per-sample values are given in Table S14.

**Figure S6:**
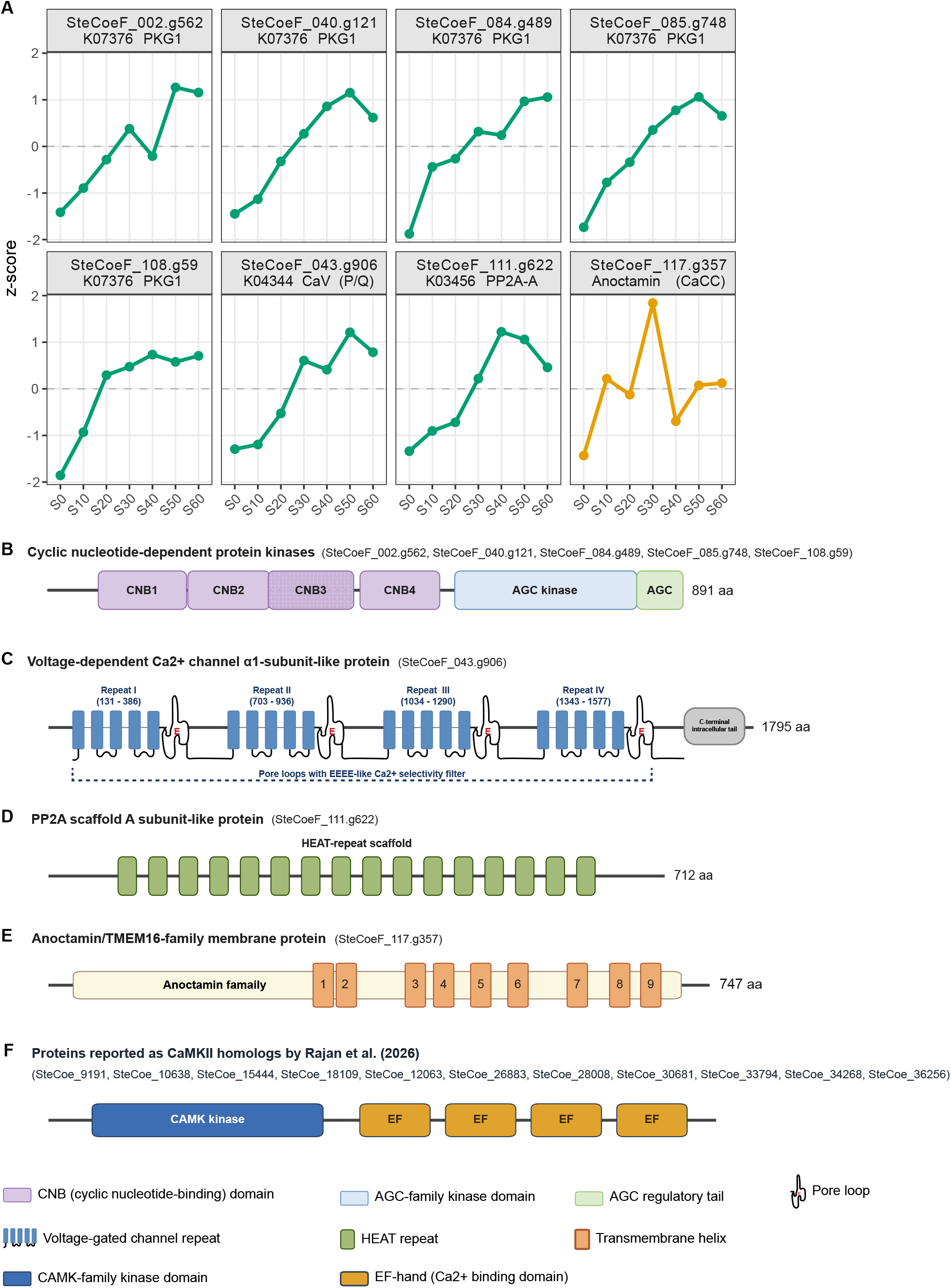
Expression trajectories and domain architectures of candidate components associated with the long-term depression signaling module. **(A)** Mean expression trajectories (row z-score across the 21 RNA-seq libraries; three biological replicates per time point) of the eight habituation-responsive genes initially assigned to the KEGG long-term depression (LTD) pathway. These include five cyclic nucleotide-dependent protein kinases (SteCoeF 002.g562, Ste-CoeF 040.g121, SteCoeF 084.g489, SteCoeF 085.g748, and SteCoeF 108.g59), one voltage-gated calcium channel α1-subunit-like protein (SteCoeF 043.g906), one protein phosphatase 2A (PP2A) scaffold A subunit-like protein (SteCoeF 111.g622), and one anoctamin/TMEM16-family membrane protein (SteCoeF 117.g357). **(B–E)** Conserved domain architectures of the curated LTD-pathway components retained for downstream analysis. **(B)** PKG paralogs possess the characteristic tandem cyclic nucleotide-binding (CNB) domains followed by an AGC-family kinase domain and regulatory tail. **(C)** The voltage-gated calcium channel α1-subunit-like protein contains four homologous channel repeats with conserved pore-loop regions. **(D)** The PP2A scaffold A subunit-like protein contains canonical HEAT-repeat domains. **(E)** The anoctamin/TMEM16-family protein contains the conserved anoctamin domain and nine predicted transmembrane helices but lacks features characteristic of the canonical LTD signaling module and was therefore excluded from the curated pathway components. **(F)** Domain organization of proteins reported as CaMKII homologs by Rajan et al. (2026). All candidates contain a CAMK-family kinase domain followed by multiple EF-hand calcium-binding domains but lack the CaMKII association domain required for canonical metazoan CaMKII holoenzyme assembly. Their domain architecture is therefore more consistent with CDPK/CDPK-like kinases than canonical animal CaMKII. Domain annotations are based on HMMER/Pfam and InterPro analyses. Source data are provided in Table S18.

